# An ATM-PPM1D Circuit Controls the Processing and Restart of DNA Replication Forks

**DOI:** 10.1101/2025.05.13.652823

**Authors:** Yiting Cao, Yingzheng Wang, Jumana Badar, Khoula Jaber, Vitor Marcel Faça, Tyler C. Huang, Eliz Çelik, Marcus B. Smolka

**Author notes:** These authors contributed equally to this work (Yiting Cao, Yingzheng Wang, Jumana Badar). To whom correspondence should be addressed: Marcus B. Smolka.

## Abstract

In response to DNA replication stress, DNA damage signaling kinases inhibit origin firing and promote the remodeling and stabilization of replication forks, leading to a systemic reduction in DNA synthesis, which collectively protects genomic integrity. Little is understood about the regulatory mechanisms of replication stress recovery, including the mechanisms involved in the restart of stalled replication forks. Here, we identify the oncogenic phosphatase PPM1D/WIP1 as a regulator of replication fork restart. During recovery from replication stress, PPM1D prevents excessive MRE11- and DNA2-dependent nucleolytic degradation of stalled forks. Loss of PPM1D function leads to defects in RAD51 recruitment to chromatin and impairs RAD51-dependent fork restart. Phosphoproteomic analysis reveals that PPM1D regulates a network of ATM substrates, several of which are phosphorylated at an S/T-Q-(E/D)^n^ motif. Strikingly, inhibition of ATM suppresses the deleterious consequences of impaired PPM1D function at replication forks, enabling timely fork restart. The dominant effect of ATM hyper-signaling in suppressing fork restart occurs, in part, through the excessive engagement of 53BP1 and consequent RAD51 antagonization. These findings uncover a new mode of ATM signaling responding to fork stalling and highlight the need for PPM1D to restrain ATM signaling and facilitate proper fork restart.

## INTRODUCTION

DNA replication occurs within a highly complex and dynamic genomic landscape. The progression of DNA replication forks is often slowed or impeded by various obstacles, such as DNA lesions or adducts, secondary DNA structures, transcriptional intermediates, compacted chromatin, and the shortage of nucleotides and other essential replication factors (*1*, *2*). When replication forks stall, they become vulnerable structures prone to degradation by nucleases, which can lead to the extensive single-stranded DNA (ssDNA) accumulation and/or the formation of double-stranded breaks (DSBs) (*3*). The ability of cells to maintain the integrity of stalled replication forks largely relies on the ATR kinase, a member of the phosphatidylinositol 3-kinase–related kinase (PIKK) family, which senses the exposure of ssDNA at replication forks (*4*). Upon fork stalling, ATR and its downstream effector CHK1 suppress late origin firing to prevent the exhaustion of key replication factors, including nucleotides, replication protein A (RPA), and the RFC/PCNA complex (*5*, *6*). In parallel, ATR signaling restrains the activity of nucleases, such as MRE11, EXO1, and MUS81, to limit excessive fork processing (*7–9*). Cells also rely on structural remodeling of replication forks to avoid nucleolytic degradation (*10*). For instance, a replication fork regresses into a four-way DNA junction, known as fork reversal, allowing nascent DNA strands to be stabilized by protective factors such as RAD51, BRCA1, and BRCA2 (*3*, *11*). This reversal process is catalyzed by DNA translocases, including SMARCAL1, HLTF, and ZRANB3 (*12*), and is tightly regulated by the DNA damage response kinases ATR and DNA-PKcs (*13*, *14*).

While fork reversal and protection are important for stabilizing replication forks, the restart of stalled replication forks is essential for the timely completion of DNA replication and the preservation of genomic integrity (*15*). Stalled forks that undergo regression can be restarted by branch migration, via the RECQ1 helicase, which is considered the primary choice for fork restart given the minimal chance for error-prone DNA processing (*16*). Alternatively, reversed forks can be restarted through regulated processing of the regressed ends by helicases and nucleases, such as WRN and DNA2, enabling RAD51-mediated homologous recombination (HR) transactions for restoring a functional replication fork (*17*). In addition, forks that collapse may undergo HR-mediated repair transactions to restore a functional fork, albeit with an increased risk of initiating mutagenic processes such as break-induced replication (BIR) (*18*). Engagement of NHEJ-mediated repair into fork-associated DSBs also has the potential of generating deleterious outcomes, such as chromosomal translocations, especially when single-ended DSBs are generated (*19*, *20*). Moreover, processes that prevent the stalling of a replication fork, such as PrimPol-mediated repriming or translesion synthesis, can mitigate the potential deleterious outcomes associated with stalled or collapsed forks, although they also introduce mutations and other genomic instabilities (*21*). Overall, the ability of cells to restart replication forks in a regulated and error-free manner is crucial to protect genome integrity upon replication stress. Since replication fork restart mechanisms need to counteract the mechanisms that stabilize stalled forks, including fork reversal and inhibition of nucleolytic processing, cells must tightly control the balance of fork-stabilizing and fork-restarting mechanisms during replication stress recovery.

While ATR is the central kinase in the response to stalled forks, the other PIKKs, ATM and DNA-PKcs, were also reported to engage in the response to various forms of replication stress. For example, in response to replication stress induced by the B-family DNA polymerase inhibitor aphidicolin, ATM was shown to phosphorylate several proteins, including H2AX, Claspin, BRCA1, MCM3, MCM6 and RIF1 (*22*), in a manner dependent on RAD18, WRNIP1, and ATMIN (*23*). Consistent with its major role in the double-strand break (DSB) response, ATM is robustly activated by agents that cause replication fork-associated DSBs, such as the topoisomerase inhibitors camptothecin and etoposide (*24*, *25*). Differences in the cellular response to stalled versus broken replication forks have been characterized at the proteomic level, which revealed a role for ATM in promoting HR-mediated repair specifically at broken forks by impairing recruitment of the anti-resection BRCA1-A complex (*25*). DNA-PKcs, which also has canonical roles in the DSB response, was shown to be recruited to both ongoing and stalled replication forks, where it mediates drug-induced replication fork reversal and promotes fork stability through mechanisms that remain unknown (*14*, *26–28*).

Since PIKK signaling halts core cellular processes such as DNA replication and cell cycle progression (*29*, *30*), mechanisms for the deactivation of PIKK signaling must be in place to enable proper recovery from replication stress. Multiple phosphatases have been implicated in antagonizing PIKK signaling, including PP1, PP2A, PPM1G and PPM1D (*31–33*). The phosphatase PPM1D (protein phosphatase Mn^2+^/Mg^2+^-dependent 1D) has been shown to play prominent roles in dephosphorylating and antagonizing ATM, p53 and CHK2 (*34–37*). PPM1D is overexpressed in multiple tumor types, including breast, ovarian, and glioblastoma tumors (*38*), which can occur through gene amplification or truncating mutations that remove a degron at its C-terminal domain (*39*, *40*). Elevated PPM1D protein levels counteract DNA damage signaling and are proposed to play a major role in reducing p53-mediated apoptosis (*39*, *41*, *42*). Not surprisingly, PPM1D overexpression has been implicated in chemotherapy resistance (*39*, *43*).

Here we find that PPM1D/WIP1 is required for the restart of stalled replication forks. Upon recovery from replication stress, PPM1D prevents the excessive nucleolytic degradation of stalled forks. Loss of PPM1D function leads to defects in RAD51 recruitment to chromatin and impairs RAD51-dependent fork restart. Phosphoproteomic analysis reveals that PPM1D regulates a network of proteins involved in ATM signaling, several of which are targeted by ATM at an S/T-Q-(E/D)^n^ motif. Strikingly, inhibition of ATM suppresses the deleterious consequences of impaired PPM1D function at replication forks, enabling timely fork restart. The dominant effect of ATM hyper-signaling in suppressing fork restart occurs through the excessive engagement of 53BP1 and consequent RAD51 antagonization. These findings uncover a new mode of ATM signaling responding to fork stalling and highlight the need for PPM1D to limit ATM signaling and enable proper fork restart.

## RESULTS

### PPM1D is required for proper restart of stalled replication forks

In search for phosphatases required for proper fork restart, we focused on the oncogenic phosphatase PPM1D (PP2Cδ/WIP1), since it was previously shown to antagonize PIKK signaling (*34*, *36*, *44–47*). PPM1D is a monomeric phosphatase for which a potent and selective inhibitor is available (GSK2830371; herein referred to as PPM1Di) (*48*), which allowed us to assess the importance of PPM1D during recovery from replication stress. U2OS cells were treated with hydroxyurea (HU) for 30 minutes, a condition previously reported to promote replication fork stalling without causing fork collapse (*49*). Cells were then washed and released in the presence or absence of PPM1Di, and DNA synthesis was assessed by monitoring the incorporation of the nucleotide analog EdU (Fig. 1a). As shown in Figs. 1a-b, PPM1Di reduced the ability of cells to incorporate EdU, suggesting a defect in replication fork restart and/or origin firing. A similar result was observed in PPM1D-knockout U2OS cells, supporting that the observed effects were not due to off-target inhibitor activity (Figs. 1c and S1a-b). DNA combing assay in U2OS cells revealed that the reduction in EdU incorporation caused by PPM1Di was associated with an increase in stalled forks and a reduction in fork restart (Figs. 1d and S1c). Consistent results were observed using DNA fiber assays in both U2OS and HCT116 cells (Fig. S1f). Additionally, both knockdown and knockout of PPM1D significantly increased the frequency of stalled forks. Notably, in PPM1D knockout cells, siPPM1D neither increases fork stalling nor decreases fork restart, supporting the specificity of the siPPM1D-mediated depletion (Figs. 1e and S1d). PPM1Di did not reduce origin firing (Fig. S1f), consistent with a defect in fork restart being the major cause of the observed reduction in DNA synthesis during recovery from replication stress. This notion is also supported by the additive effect of PPM1Di in combination with the CDC7 inhibitor, an inhibitor of origin firing, in reducing DNA synthesis (Fig. S1g). Moreover, the overall abundance of PPM1D is increased in S-phase cells (Figs. 1f-g), consistent with previously published results (*50*), and supporting a potential role for PPM1D in DNA replication. Taken together, these results reveal a role for PPM1D in the control of replication fork restart.

**Figure 1.**
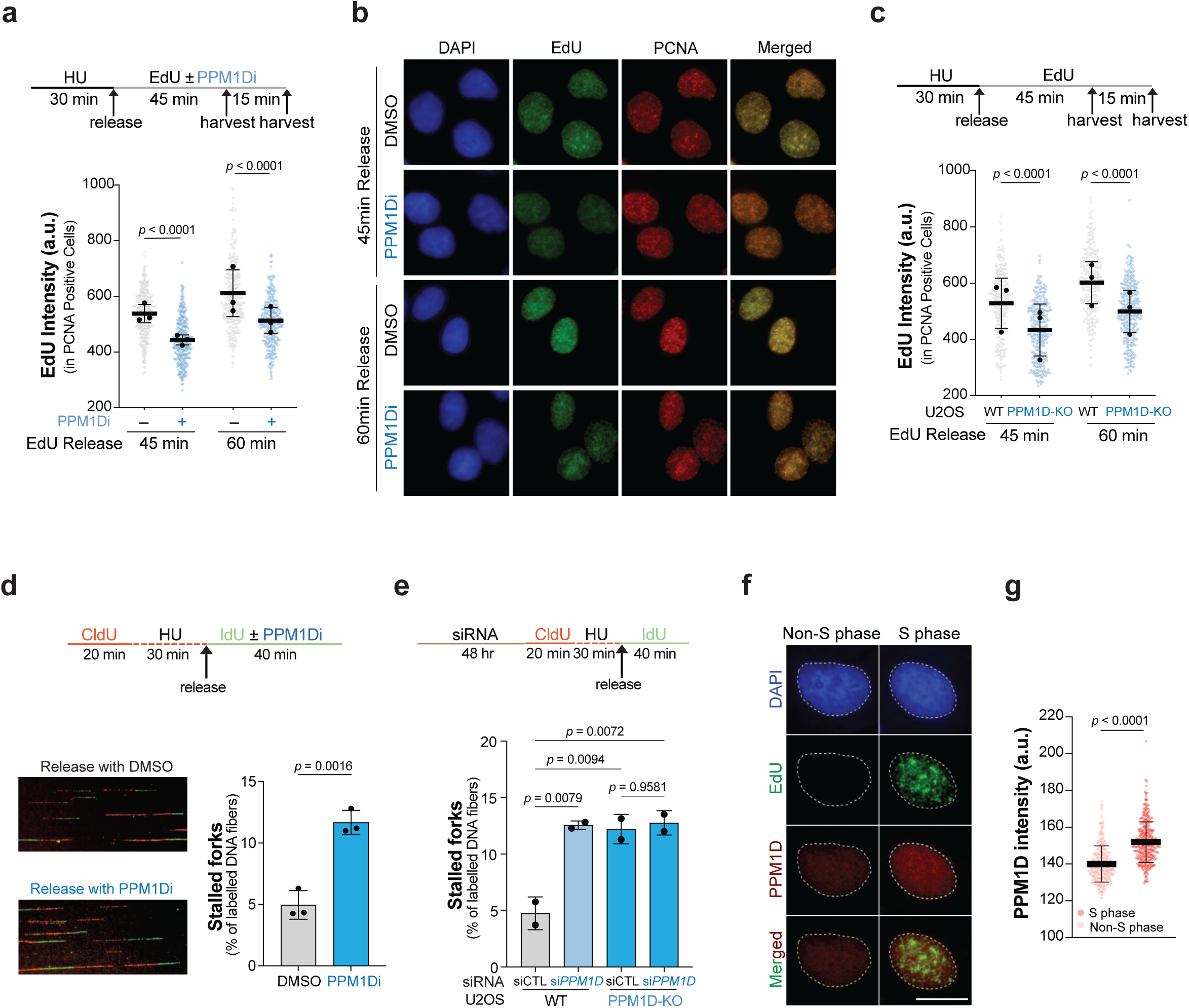
PPM1D regulates replication fork restart. (a) EdU incorporation analysis in U2OS cells. Top: experimental scheme illustrating treatment with the PPM1D inhibitor (PPM1Di; GSK2830371; 10 μM). Bottom: dot plot showing EdU intensity in PCNA-positive cells pooled from three independent biological replicates (at least 104 cells per condition per replicate). The mean EdU intensity from each replicate is indicated by black dots. *p*-values were determined by one-way ANOVA with Tukey’s multiple comparisons test. Error bars represent standard deviation. (b) Representative immunofluorescence images from the EdU incorporation assay shown in (a). Scale bar, 10 μm. (c) EdU incorporation analysis in U2OS wild-type (WT) and PPM1D-knockout (KO) cells. Top: experimental scheme. Bottom: dot plot showing EdU intensity in PCNA-positive cells pooled from three independent biological replicates (at least 83 cells per condition per replicate). The mean EdU intensity from each replicate is indicated by black dots. *p*-values were determined by one-way ANOVA with Tukey’s multiple comparisons test. Error bars represent standard deviation. (d) DNA combing assay in U2OS cells. Top: experimental scheme. Bottom left: Representative images of DNA combing assay. Bottom right: bar graph showing the percentages of stalled replication forks in cells treated with or without PPM1Di (10 μM). A minimum of 2,720 fibers were quantified per condition from three independent biological replicates. *p*-values were calculated using Student’s t-test. Error bars represent standard deviation. (e) DNA combing assay in U2OS cells transfected with control and PPM1D siRNA (siCTL and si*PPM1D*). Top: experimental scheme. Bottom: bar graph showing the percentages of stalled replication forks in U2OS wild-type (WT) and PPM1D-knockout (KO) cells. A minimum of 1,644 fibers were quantified per condition from two independent biological replicates. *p*-values were calculated using one-way ANOVA with Tukey’s multiple comparisons test. Error bars represent standard deviation. (f) Representative immunofluorescence images showing co-staining of PPM1D (red) and EdU (green) detected via anti-PPM1D antibody and click chemistry, respectively. Scale bar, 10 μm. (g) Dot plot showing the PPM1D intensity in EdU-negative and EdU-positive U2OS cells. At least 328 cells were analyzed per condition. *p*-value was calculated using Student’s t-test. Error bars represent standard deviation.

### PPM1D prevents the nucleolytic processing of stalled replication forks during replication stress recovery

To gain insights into how loss of PPM1D function impairs replication forks during replication stress recovery, we monitored RPA32 foci as a proxy for ssDNA accumulation. U2OS cells were treated with HU for 30 minutes and released in the presence or absence of PPM1D function. As shown in Figs. 2a-d, both PPM1Di and PPM1D knockout caused an increase in RPA32 foci in PCNA-positive cells. We also monitored the accumulation of ssDNA at nascent DNA strands during release from HU arrest using BrdU labeling followed by non-denaturing immunofluorescence and observed a significant increase in nascent ssDNA exposure in cells treated with PPM1Di (Figs. 2e and S2a-b). We interpret the increased BrdU signal on nascent DNA strands as likely originating from ssDNA at regressed replication forks. Consistent with this notion, cells lacking SMARCAL1, a translocase involved in fork reversal, did not exhibit significantly increased levels of RPA32 foci or stalled forks upon PPM1Di treatment during HU release (Figs. 2f-g and S2c). Moreover, inhibition of the nucleases MRE11 or DNA2 suppressed RPA32 foci accumulation (Figs. 2h-i), supporting the model that, upon PPM1Di treatment during HU release, the action of these nucleases generates excessive ssDNA at stalled replication forks. Notably, DNA2 seemed to play a more prominent role, with DNA2i completely rescuing the effect of PPM1Di, consistent with previous reports showing that this is the key nuclease involved in the hyper-processing of reversed replication forks (*17*). Overall, these results point to a role for PPM1D in preventing the aberrant nucleolytic processing of stalled replication forks during recovery from replication stress.

**Figure 2.**
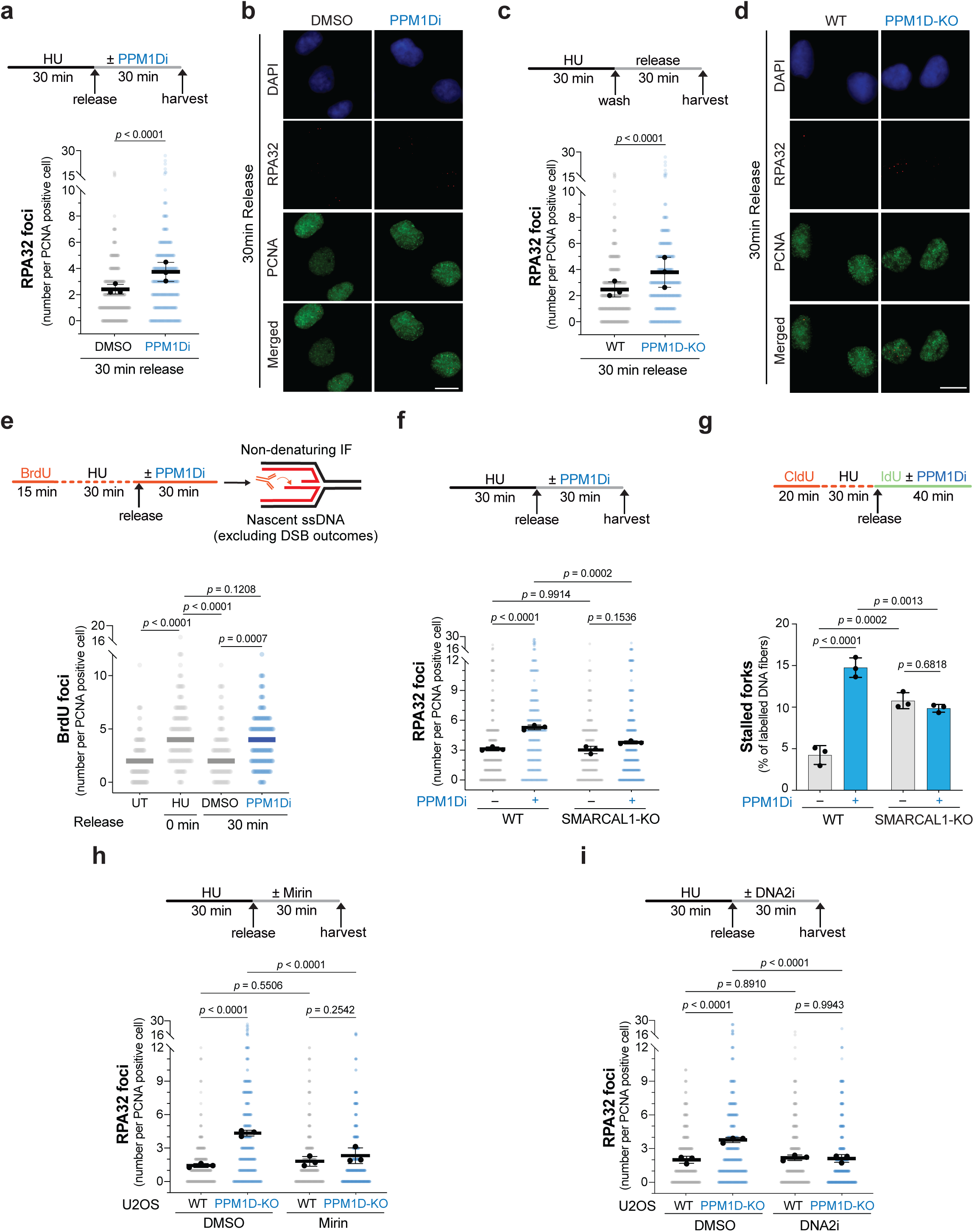
PPM1D counteracts nucleolytic processing of stalled replication forks. (a) RPA32 foci analysis in U2OS cells. Top: experimental scheme illustrating treatment with the PPM1D inhibitor (PPM1Di, 10 µM). Bottom: dot plot showing the number of RPA32 foci in PCNA-positive cells pooled from three independent biological replicates (at least 101 cells per condition per replicate). The mean number of RPA32 foci from each replicate is indicated by black dots. *p*-value was calculated using Student’s t-test. Error bars represent standard deviation. (b) Representative immunofluorescence images corresponding to (a). Scale bar, 10 μm. (c) RPA32 foci analysis in U2OS wild-type (WT) and PPM1D-knockout (KO) cells. Top: experimental scheme. Bottom: dot plot shows the number of RPA32 foci in PCNA-positive cells pooled from three independent biological replicates (at least 101 cells per condition per replicate). The mean number of RPA32 foci from each replicate is indicated by black dots. *p*-value was calculated using Student’s t-test. (d) Representative immunofluorescence images corresponding to (c). Scale bar, 10 μm. (e) Analysis of nascent single-stranded DNA (ssDNA) in U2OS cells. Top: experimental scheme showing BrdU immunostaining under non-denaturing conditions following the indicated treatment. Bottom: dot plot shows the number of BrdU foci in PCNA-positive cells from one representative experiment. Two biological replicates were performed; results from the second biological replicate are shown in Fig. S2b. *p*-values were calculated using one-way ANOVA. (f) RPA32 foci analysis in U2OS WT and SMARCAL1-KO cells. Top: experimental scheme illustrating treatment with the PPM1Di (10 μM). Bottom: dot plot shows the number of RPA32 foci in PCNA-positive cells pooled from three independent biological replicates (at least 73 cells per condition per replicate). The mean number of RPA32 foci from each replicate is indicated by black dots. *p*-values were calculated using one-way ANOVA with Tukey’s multiple comparisons test. (g) DNA fiber assay in U2OS WT and SMARCAL1-KO cells. Top: experimental scheme. Bottom: bar graph shows the percentage of stalled replication forks in the presence or absence of PPM1Di (10 μM). At least 942 fibers per condition were quantified across three independent biological replicates. *p*-values were calculated using one-way ANOVA with Tukey’s multiple comparisons test. Error bars represent standard deviation. (h-i) RPA32 foci analysis in U2OS WT and PPM1D-KO cells treated with or without MRE11 inhibitor (Mirin; 10 μM) (h) or DNA2 inhibitor (DNA2i; 20 μM) (i), as shown in the experimental schemes. Dot plots show the number of RPA32 foci in PCNA-positive cells pooled from three independent biological replicates (at least 97 (h) or 99 (i) cells per condition per replicate). The mean number of RPA32 foci from each replicate is indicated by black dots. *p*-values were calculated using one-way ANOVA with Tukey’s multiple comparisons test. Error bars represent standard deviation.

### PPM1D antagonizes ATM signaling during recovery from fork stalling

To delineate the contribution of PPM1D to signaling downregulation during recovery from replication stress, we compared the phosphoproteomes of mock-treated versus PPM1Di-treated cells during release from HU arrest. The generated dataset was crossed with data from a separate analysis comparing the phosphoproteomes of cells arrested by HU with the phosphoproteome of cells that were HU-arrested and then released, yielding a dataset of 10,833 overlapping phosphosites identified in both phosphoproteomic experiments (Fig. 3a, Table S1). PPM1Di caused extensive changes in signaling during replication stress recovery, resulting in at least a two-fold increase in the abundance of about 18% of the phosphorylation events found to be downregulated during release from HU arrest (Figs. 3a-b). While this finding defines PPM1D as a prominent phosphatase acting during replication stress recovery, it also implies that additional phosphatases must contribute to the dephosphorylation of other signaling events downregulated during replication stress recovery. Notably, out of the phosphorylation events upregulated by at least 2-fold upon treatment with PPM1Di, 22.5% contained a phosphorylation event at the SQ/TQ consensus motif (Fig. 3c), suggesting that PPM1D displays some level of selectivity for antagonizing PIKK signaling. Consistent with previous reports, we found that PPM1D displays a preference for targeting phosphorylation at an S/T-Q-(E/D)^n^ motif (Fig. 3d) (*47*, *51*). Notably, the phosphorylation events at S/T-Q-(E/D)^n^ motifs most highly regulated by PPM1D included several known substrates of the ATM kinase, such as RAP80, NUMA1, SMC1A, and 53BP1 (*52–55*), suggesting that PPM1D plays a prominent role in antagonizing ATM signaling. We confirmed this prediction by monitoring the phosphorylation of KAP1 at serine 824, a well-established ATM substrate within the S/T-Q-(E/D)^n^ motif (*56*, *57*). As shown in Figs. 3e-g, while PPM1D inhibition during release from HU treatment caused a drastic increase in KAP1 phosphorylation, it led to a more subtle increase in CHK1 phosphorylation, consistent with the model that PPM1D plays a more prominent role in antagonizing ATM signaling compared to ATR signaling. Notably, we did not observe changes in the level of ATM autophosphorylation (Fig. 3h), suggesting that PPM1D acts on ATM downstream substrates as opposed to acting on ATM itself. Overall, our results establish PPM1D as a key phosphatase antagonizing ATM signaling during replication stress recovery.

**Figure 3.**
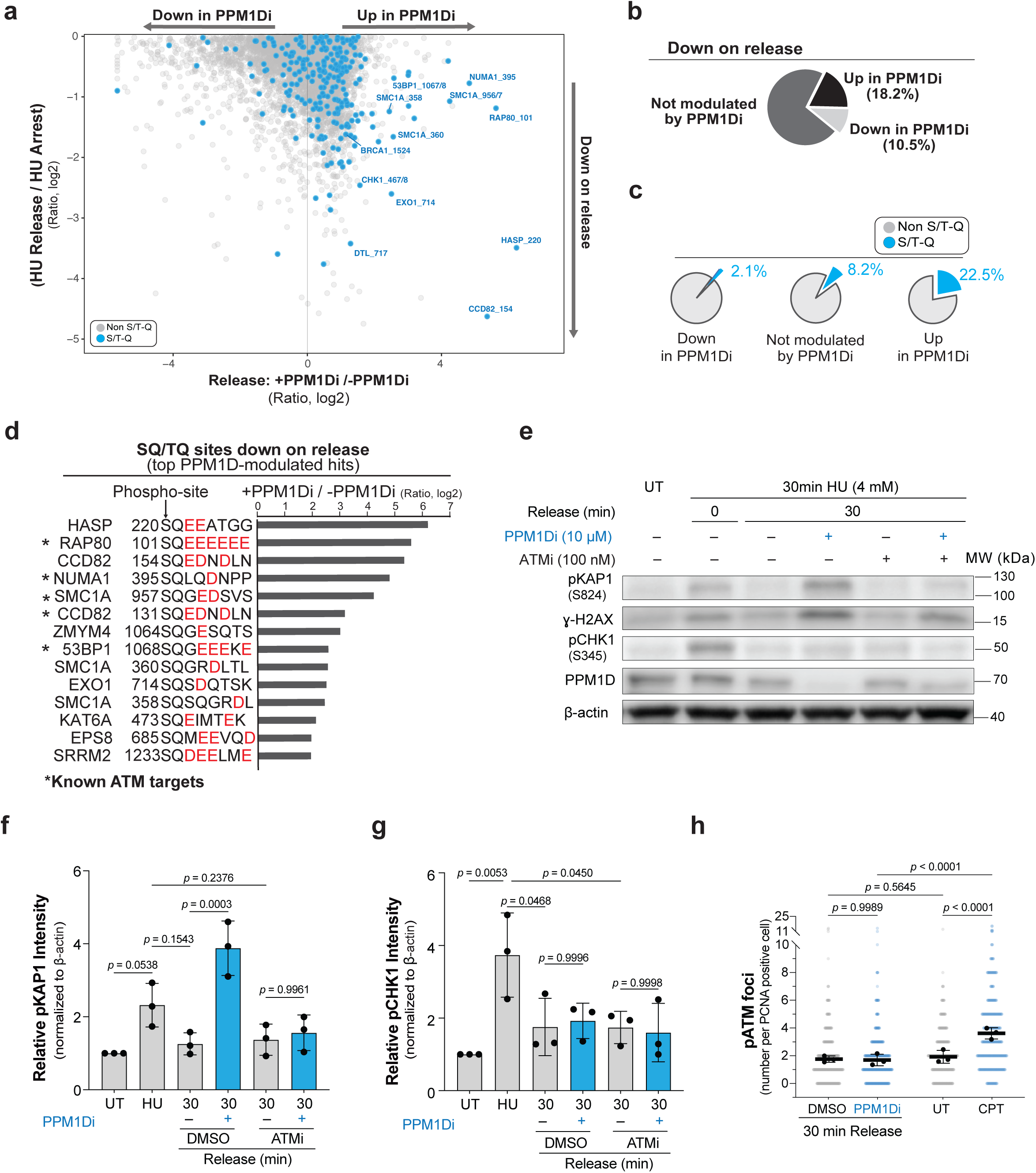
PPM1D antagonizes ATM signaling during replication stress recovery. (a) Scatter plot showing all phosphosites downregulated during replication stress recovery. Phosphosites from the HU Release/HU Arrest comparison are categorized along the *y*-axis as either PPM1D-independent (left and center) or PPM1D-dependent (right). (b) Pie chart summarizing all phosphosites downregulated during replication stress recovery, illustrating the proportions of PPM1D-independent and PPM1D-dependent sites. (c) Pie chart highlighting the subset of S/T-Q phosphosites within the PPM1D-independent and PPM1D-dependent groups shown in (b). (d) Bar graph demonstrating the enrichment of the consensus phospho-motifs (S/T-Q or S/T-Q-(D/E)^2+^) in cells treated with or without PPM1D inhibitor (PPM1Di). (e) Immunoblot analysis of U2OS cells treated with hydroxyurea (HU; 4 mM), PPM1Di (10 µM), and/or ATMi (100 nM) for the indicated times. (f-g) Quantification of relative protein levels of pKAP1 (f) and pCHK1 (g) based on immunoblot analysis in (e). Bar graphs show normalized protein expression levels relative to β-actin across three independent biological replicates. The normalized protein expression level from each replicate is indicated by black dots. *p*-values were calculated using one-way ANOVA with Tukey’s multiple comparisons test. Error bars represent standard deviation. (h) Phospho-ATM (pATM, S1981) foci analysis in U2OS cells treated with or without PPM1Di. Dot plot shows the number of pATM foci in PCNA-positive cells pooled from three independent biological replicates (at least 91 cells per condition per replicate). The mean number of pATM foci from each replicate is indicated by black dots. *p*-values were calculated using one-way ANOVA with Tukey’s multiple comparisons test. Error bars represent standard deviation.

### Unrestrained ATM signaling causes aberrant fork processing and fork restart defects upon loss of PPM1D function

Since inhibition of PPM1D resulted in the increased phosphorylation of multiple ATM substrates during replication stress recovery, we next tested whether inhibition of ATM could suppress the defects we observed with fork restart and nucleolytic processing. PPM1Di alone did not cause a strong reduction in EdU incorporation when compared to its effect during replication stress release (Fig. S3a). Strikingly, inhibition of ATM during release from HU treatment fully rescued the increased accumulation of RPA32 foci (Figs. 4a-b) and restored EdU incorporation in PPM1Di-treated cells (Figs. 4c-d and S3a). ATMi also rescued the accumulation of stalled forks and fork degradation caused by PPM1Di, as monitored by DNA fiber assays (Figs. 4e-f and S3b-c). Furthermore, inhibition of ATM, but not ATR, was able to rescue the proliferation defect caused by PPM1Di (Fig. 4g), reinforcing the hypothesis that PPM1D plays a key role in selectively antagonizing ATM-dependent signaling. Taken together, these results support a model in which PPM1D dephosphorylates ATM substrates during recovery from replication stress, thereby preventing their toxic accumulation and the consequent hyperactivation of nucleases whose action results in ssDNA accumulation and defects in fork restart.

**Figure 4.**
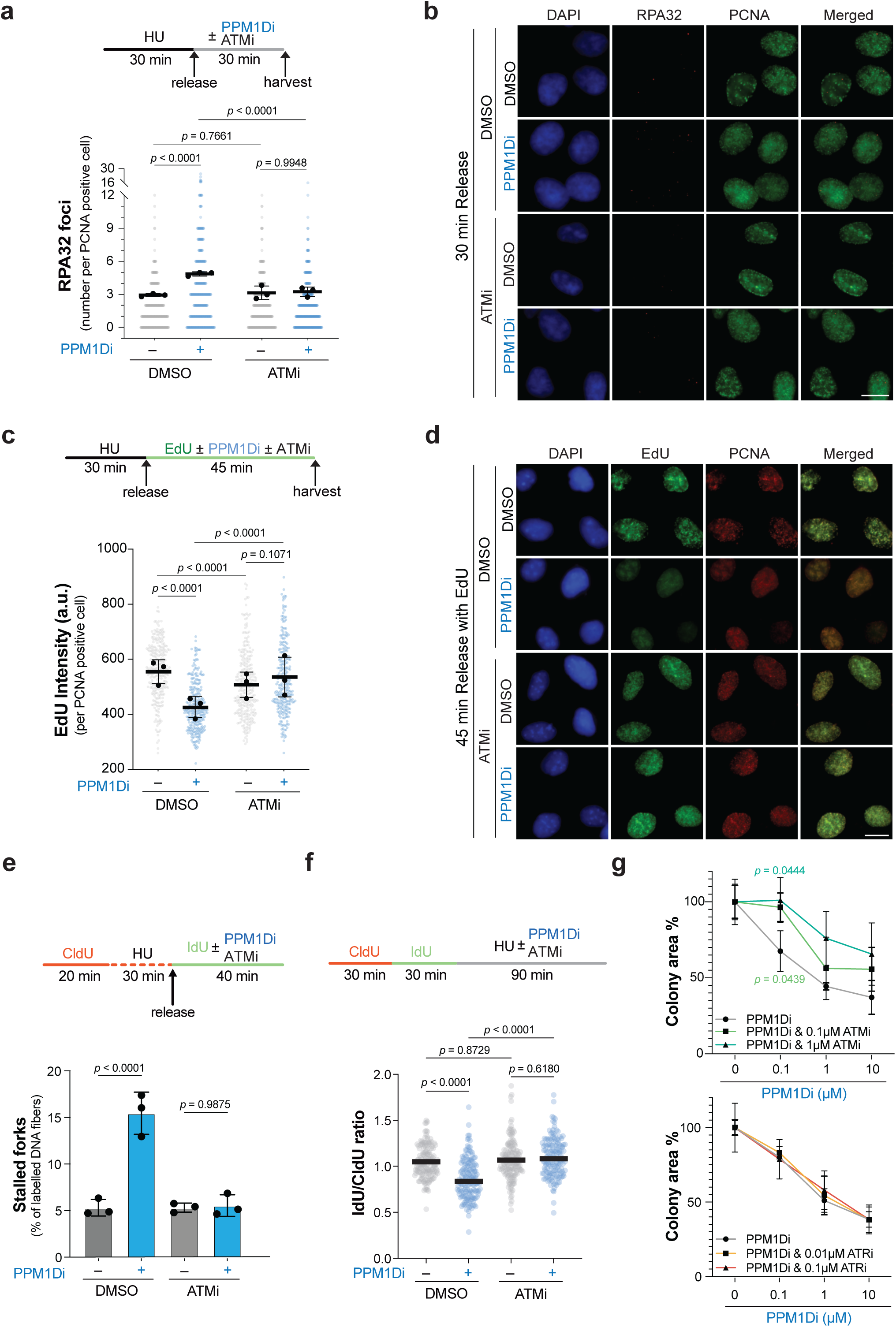
ATM inhibition rescues defects in replication fork processing and restart caused by loss of PPM1D function. (a) RPA32 foci analysis in U2OS cells. Top: experimental scheme showing treatment with PPM1Di and ATM inhibitor (ATMi; AZD0156; 100 nM). Bottom: dot plot shows the number of RPA32 foci in PCNA-positive cells pooled from three independent biological replicates (at least 93 cells per condition per replicate). The mean number of RPA32 foci from each replicate is indicated by black dots. *p*-values were calculated using one-way ANOVA with Tukey’s multiple comparisons test. Error bars represent standard deviation. (b) Representative immunofluorescence images corresponding to the RPA32 foci analysis shown in (a). Scale bar, 10 μm. (c) EdU incorporation analysis in U2OS cells. Top: experimental scheme showing treatment with PPM1Di and ATMi. Bottom: dot plot shows the EdU intensity in PCNA-positive cells pooled from three independent biological replicates (at least 95 cells per condition per replicate). The mean EdU intensity from each replicate is indicated by black dots. *p*-values were determined by one-way ANOVA with Tukey’s multiple comparisons test. Error bars represent standard deviation. (d) Representative immunofluorescence images from the EdU incorporation assay shown in (c). Scale bar, 10 μm. (e) DNA fiber assay in U2OS cells. Top: experimental scheme showing treatment with PPM1Di and ATMi, as indicated. Bottom: bar graph shows the percentage of stalled replication forks under each treatment condition. A minimum of 2,929 fibers per condition were analyzed from three independent biological replicates. *p*-values were calculated using one-way ANOVA. Error bars represent standard deviation. (f) DNA protection assay in U2OS cells. Top: experimental scheme. Bottom: dot plot showing the ratio of IdU/CldU tract lengths as an indicator of replication fork degradation in one representative experiment. At least 150 fibers were analyzed per condition. Results from the second biological replicate are shown in Fig. S3c. *p*-values were calculated using one-way ANOVA. (g) Clonogenic survival assay of U2OS cells co-treated with ATMi, ATRi, and PPM1Di. Colony area was normalized to the respective untreated PPM1Di control group for each treatment condition. Data represent three independent biological replicates. *p*-values were assessed using Student’s t-test.

### PPM1D controls RAD51 chromatin recruitment and RAD51-mediated fork restart

To better characterize the effects of unrestrained ATM signaling during replication stress recovery, we monitored RAD51, a recombinase protein with separable roles in protecting reversed forks from degradation and mediating fork restart via homologous recombination (*58*). Immunofluorescence-based analysis revealed that the number of RAD51 foci was significantly reduced in cells treated with PPM1Di during recovery from replication stress (Figs. 5a-b). This finding is consistent with a recent report showing that PPM1D inhibition suppresses RAD51 foci formation in DIPG cells, as well as in U2OS cells following exposure to ionizing radiation (*59*, *60*). In addition, chromatin fractionation also revealed a reduction of RAD51 in the chromatin-bound fraction upon PPM1D inhibition during recovery from replication stress (Fig. 5c). We hypothesized that PPM1D promotes RAD51-dependent fork restart during replication stress recovery. To test this, we performed the DNA fiber assay to monitor persistent fork stalling in cells treated with the RAD51 inhibitor B02, in the presence or absence of PPM1Di (Figs. 5d and S4). Consistent with previous reports highlighting the importance of RAD51 for fork restart (*49*), RAD51 inhibition alone increased the incidence of stalled forks. Cells co-treated with RAD51i and PPM1Di during recovery from HU arrest did not display increased incidence of stalled forks compared to cells treated with RAD51i alone. These findings suggest that PPM1D and RAD51 function through the same pathway to facilitate fork restart. The degree of fork stalling in RAD51i-treated cells was higher than in PPM1Di-treated cells, congruent with a model whereby PPM1D functions as a regulator of RAD51 roles in ensuring proper fork restart. Such regulation is likely to involve PPM1D’s role in antagonizing ATM signaling as inhibition of ATM prevents reduction of RAD51 foci in PPM1D-KO cells (Fig. 5e). Additionally, we observed that PPM1D inhibition caused a rapid increase in 53BP1 foci during the recovery from replication stress, and that the increase was dependent on ATM (Fig. 5f). 53BP1 is a bona-fide ATM target (*61*, *62*) and one of the top PPM1D-regulated targets identified in our phosphoproteomic screen (Fig. 3d). Recent studies have shown that 53BP1 counteracts RAD51 loading at replication forks independently of its role in antagonizing resection (*63*). Therefore, we reasoned that the hyperphosphorylation of 53BP1 upon PPM1D inhibition causes its accumulation at stalled replication forks, which in turn counteracts RAD51 engagement, thereby impairing proper fork restart. Consistent with this model, PPM1D inhibition did not cause a decrease in RAD51 engagement in 53BP1-KO cells (Fig. 5g).

**Figure 5.**
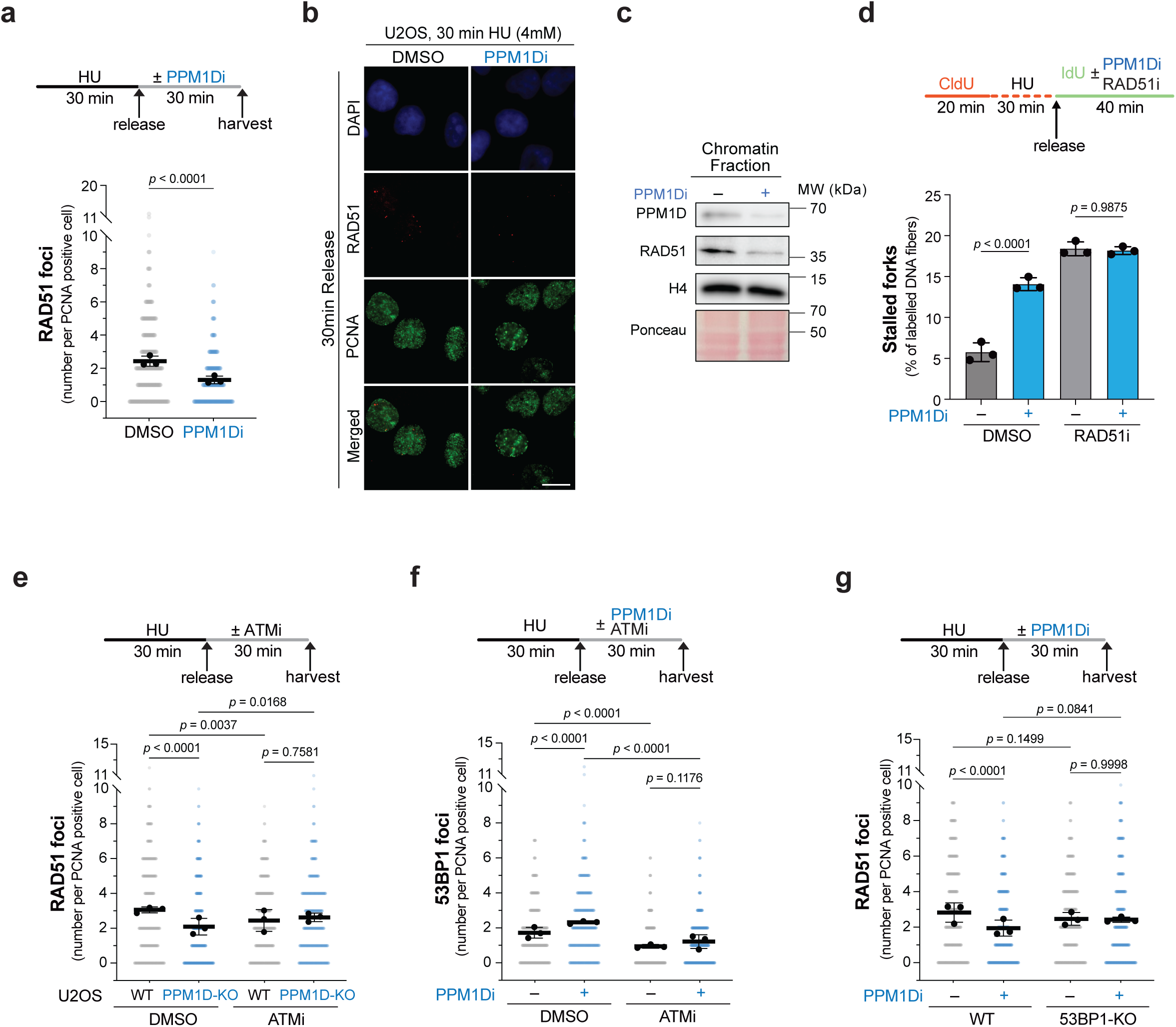
PPM1D regulates RAD51-mediated replication fork restart. (a) RAD51 foci analysis in U2OS cells. Top: experimental scheme. Bottom: dot plot shows the number of RAD51 foci in PCNA-positive cells pooled from three independent biological replicates (at least 96 cells per condition per replicate). The mean number of RAD51 foci from each replicate is indicated by black dots. *p*-value was calculated using Student’s t-test. Error bars represent standard deviation. (b) Representative immunofluorescence images corresponding to (a). Scale bar, 10 μm. (c) Immunoblot analysis of PPM1D, RAD51, and histone H4 in chromatin fractions from U2OS cells treated with or without PPM1Di. Ponceau S staining was used as a loading and total protein normalization control. (d) DNA fiber assay in U2OS cells. Top: experimental scheme showing treatment with PPM1Di and RAD51 inhibitor (RAD51i; B02; 20 μM). Bottom: bar graph shows the percentage of stalled replication forks under each condition. A minimum of 2,959 fibers per condition were quantified from three independent biological replicates. *p*-values were calculated using one-way ANOVA. Error bars represent standard deviation. (e) RAD51 foci analysis in U2OS wild-type (WT) and PPM1D-knockout (KO) cells. Top: experimental scheme. Bottom: dot plot shows the number of RAD51 foci in PCNA-positive cells pooled from three independent biological replicates (at least 102 cells per condition per replicate). The mean number of RAD51 foci from each replicate is indicated by black dots. *p*-values were calculated using one-way ANOVA with Tukey’s multiple comparisons test. Error bars represent standard deviation. (f) 53BP1 foci analysis in U2OS cells. Top: experimental scheme. Bottom: dot plot shows the number of 53BP1 foci in PCNA-positive cells pooled from three independent biological replicates (at least 96 cells per condition per replicate). The mean number of 53BP1 foci from each replicate is indicated by black dots. *p*-values were calculated using one-way ANOVA with Tukey’s multiple comparisons test. Error bars represent standard deviation. (g) RAD51 foci analysis in U2OS WT and 53BP1-knockout (KO) cells. Top: experimental scheme. Bottom: dot plot shows the number of RAD51 foci in PCNA-positive cells pooled from three independent biological replicates (at least 88 cells per condition per replicate). The mean number of RAD51 foci from each replicate is indicated by black dots. *p*-values were calculated using one-way ANOVA with Tukey’s multiple comparisons test. Error bars represent standard deviation.

### Increased phosphorylation status of ATM targets upon loss of PPM1D is not correlated with an increase in detectable DSBs

ATM is canonically activated by DSBs (*64*). However, neutral comet assay did not detect PPM1Di-dependent increase in DSBs within 30 minutes following recovery from replication stress, a time point in which PPM1D inhibition resulted in a significant increase in the ATM-dependent phosphorylation of KAP1 at serine 824 (Figs. 6a-c). These findings suggest that ATM signaling and the observed increases in the phosphorylation status of ATM substrates are independent of DSB formation during early replication stress recovery. Our finding that the autophosphorylation of ATM is not increased during release from replication stress is also consistent with this notion (Fig. 3h). To provide further evidence that PPM1D inhibition is not causing strong induction of DSBs early (∼30 minutes) during recovery from replication stress, we engineered a ProKAS biosensor (*65*) containing two sensors that differentiate ATM signaling regulated by PPM1D (ATM-PPM1D sensor) from ATM signaling that is not controlled by PPM1D (ATM sensor) (Figs. 6d-e). As shown in Figs. 6f and S5, CPT robustly increased the phosphorylation of both sensors, consistent with the canonical role of ATM in responding to DSBs. The ATM sensor did not report increased ATM activity during recovery from replication stress, in the absence or presence of PPM1D inhibitor, further supporting the absence of DSB-induced ATM signaling under these conditions. Notably, the ATM-PPM1D sensor reported a drastic increase in ATM signaling upon PPM1D inhibition during recovery from replication stress. These findings with the biosensor are congruent with the model that PPM1D inhibition is not causing DSBs, and suggest that in the context of replication fork stalling and restart, ATM may be activated by alternative DNA structures, such as the nascent DNA ends of reversed forks.

**Figure 6.**
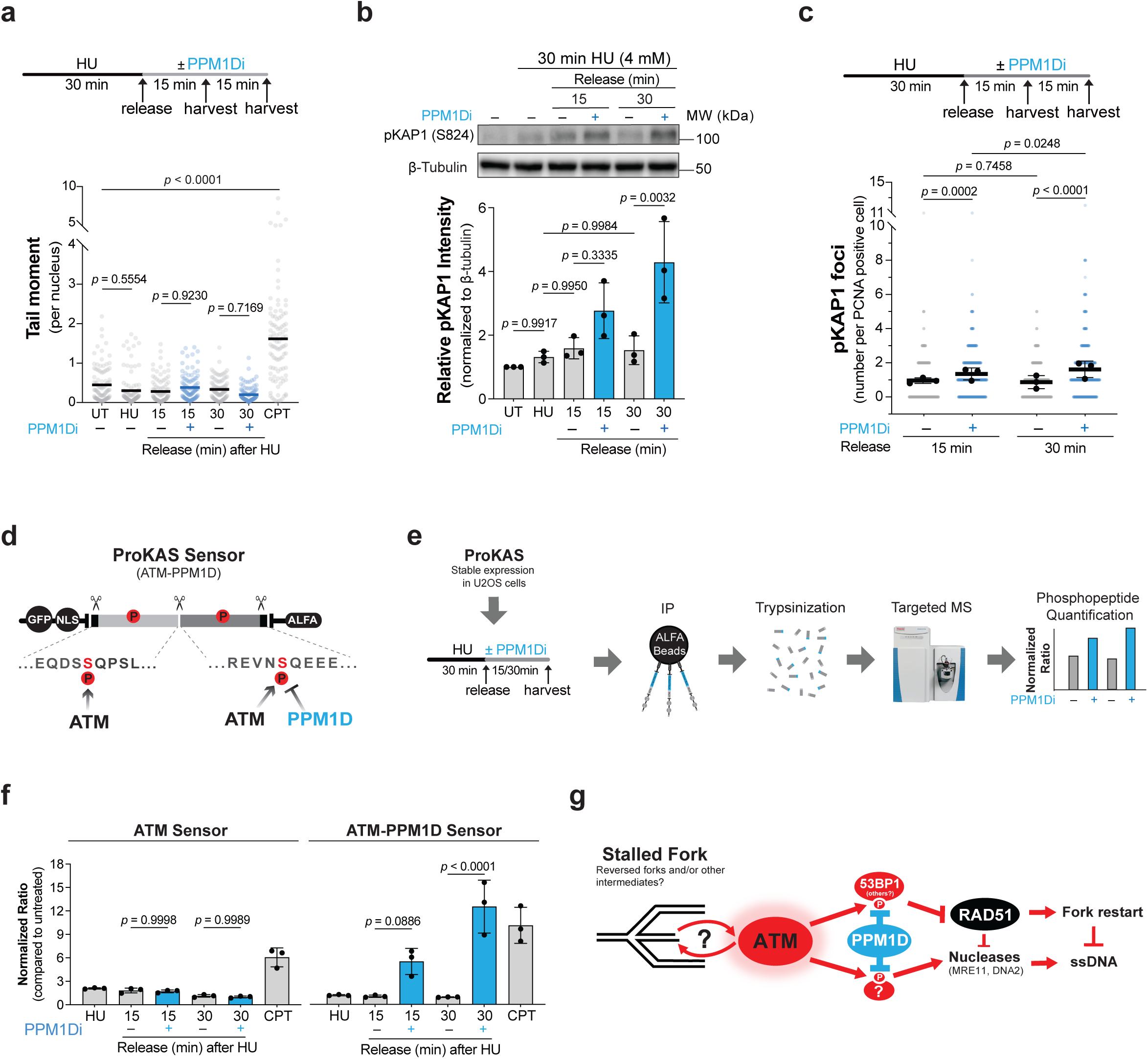
Increased ATM signaling during fork recovery in cells with impaired PPM1D function is not correlated with an increase in detectable DSB. (a) Neutral comet assay in U2OS cells. Top: experimental scheme. Bottom: dot plot shows the tail moment for each condition. At least 102 cells were quantified per condition. Treatment with 1 μM camptothecin (CPT) for 30 minutes was used as a positive control. *p*-values were calculated using one-way ANOVA. (b) Immunoblot analysis of phosphorylated KAP1 (pKAP1) in U2OS cells treated with HU (4 mM) and/or PPM1Di (10 µM) for the indicated durations. The bar graph at the bottom shows the normalized pKAP1 expression levels relative to β-tubulin from three independent biological replicates. The normalized pKAP1 expression level from each replicate is indicated by black dots. *p*-values were calculated using one-way ANOVA with Tukey’s multiple comparisons test. Error bars represent standard deviation. (c) pKAP1 (S824) foci analysis in U2OS cells. Top: experimental scheme. Bottom: dot plot shows the number of pKAP1 foci in PCNA-positive cells pooled from three independent biological replicates (at least 125 cells per condition per replicate). The mean number of pKAP1 foci from each replicate is indicated by black dots. *p*-values were calculated using one-way ANOVA with Tukey’s multiple comparisons test. Error bars represent standard deviation. (d) ProKAS (Proteomic Kinase Activity Sensor) design. Peptide sequences used in the ATM sensor (PPM1D-untargeted) and ATM-PPM1D sensor (PPM1D targeted) are shown. (e) Experimental scheme using ProKAS for detecting ATM and PPM1D activities. (f) Quantification of ATM and PPM1D activity using ProKAS sensors during replication stress recovery with or without PPM1Di. Bar graphs represent the normalized ATM activity (left) and PPM1D acitivity (right) relative to untreated condition from three independent biological replicates. *p*-values were calculated using one-way ANOVA with Tukey’s multiple comparisons test. Error bars represent standard deviation. (g) Proposed model illustrating how PPM1D promotes replication fork restart and prevents ssDNA accumulation by counteracting ATM signaling at stalled replication forks.

## DISCUSSION

The ability of cells to restart stalled replication forks is crucial for the faithful completion of DNA replication under conditions of replication stress. Despite its importance, the regulatory mechanisms that govern fork restart remain incompletely understood. Here, we identify the phosphatase PPM1D as a regulator of replication fork recovery. Our findings reveal that attenuation of ATM-mediated signaling is a central function of PPM1D during replication stress resolution. In our working model (Fig. 6g), ATM is activated at stalled forks and phosphorylates substrates such as 53BP1, which in turn interferes with RAD51 loading. Persistent ATM activity antagonizes HR-mediated fork restart and also leads to fork degradation due to reduced RAD51-mediated fork protection. PPM1D counteracts the block in RAD51 loading by dephosphorylating ATM targets. Thus, the ATM–PPM1D axis acts as a dynamic rheostat that modulates ATM signaling at stalled forks to coordinate the balance between fork protection and fork restart.

A key unresolved question is how PPM1D is recruited to stalled forks. It is possible that the build-up of ATM-dependent phosphorylation events triggers PPM1D recruitment, establishing a feedback mechanism to restrain excessive ATM signaling at forks. Future studies will be needed to define the mechanisms driving PPM1D localization and whether recruitment is influenced by specific post-translational modifications, replisome-associated factors, or fork-associated DNA structures. Moreover, our results suggest that PPM1D primarily acts on ATM substrates rather than directly on ATM itself or ATR substrates. This is consistent with PPM1D’s preference for dephosphorylating S/T-Q motifs followed by acidic residues [S/T-Q-(D/E)n motifs], a motif also preferred by ATM (*66*).

While ATM canonically responds to DSBs, our work suggests that ATM can also respond to replication stress independently of DSB formation. Under conditions of PPM1D inhibition, we observe elevated ATM signaling without detectable DSBs by neutral comet assay. Additionally, ProKAS biosensor analyses show that the branch of ATM signaling regulated by PPM1D is selectively induced during replication stress recovery, while other branches remain unchanged. Although the precise DNA structures activating ATM at stalled forks remain unclear, we favor the model that ATM senses unprocessed DNA ends at reversed forks. Nevertheless, given previous reports of ATM activation in response to altered chromatin states or oxidative stress (*67*, *68*), we cannot exclude the possibility that chromatin remodeling or oxidative stress during replication stress recovery also contributes to ATM activation.

Another outstanding question is whether ATM signaling at stalled forks plays beneficial roles in maintaining genome stability or supporting replication.One possibility is that early ATM-mediated 53BP1 recruitment to stalled forks, and the consequent suppression of RAD51 loading, allows nucleases such as MRE11 and DNA2 to perform limited resection during recovery from replication stress. This limited ssDNA production is needed for subsequent HR-mediated fork restart.Therefore, PPM1D is needed later in the recovery process to attenuate ATM signaling, reduce 53BP1 engagement and increase RAD51 loading, facilitating the transition from fork processing to HR-mediated fork restart. Importantly, the ability of 53BP1 to antagonize RAD51 loading, a role for 53BP1 also recently demonstrated by others (*63*), may prevent inappropriate or premature recombination events that could cause chromosomal rearrangements. Thus, by orchestrating these dynamic regulatory steps, the ATM–PPM1D axis could ensure a timely, error-free, and balanced coordination of fork protection, processing, and HR-mediated restart. Future investigations will be required to define the spatio-temporal regulation of PPM1D and ATM action at replication forks.

Alternatively, ATM activation at stalled forks may largely reflect an unintended response to replication intermediates resembling DSBs, such as the DNA end of a regressed fork arm. In this scenario, unchecked ATM signaling may represent a maladaptive response, with PPM1D acting as a safeguard to dampen inappropriate ATM signaling and enable proper fork restart and recovery from replication stress. In the future, it will be important to better characterize the impact of PPM1D and ATM inhibition on the accumulation of replication stress-induced genomic instabilities, such as chromosomal aberrations. Detection of HR-driven chromosomal rearrangements upon ATM inhibition would point to an important role for ATM signaling in promoting HR quality control during replication stress responses. Moreover, delayed RAD51 loading and impaired HR-mediated fork restart in the absence of PPM1D may lead to increased utilization of error-prone pathways, such as non-homologous end joining (NHEJ) or microhomology-mediated end joining (MMEJ), resulting in elevated frequencies of chromosomal translocations and radial chromosomes, which can be revealed by metaphase spread analysis.

These insights have important implications for cancer biology. PPM1D is frequently overexpressed or mutated in cancers, often leading to elevated phosphatase activity. Such dysregulation may enhance cancer cell survival under endogenous or therapy-induced replication stress by facilitating fork restart and recovery (*59*). Pharmacological targeting of PPM1D could synergize with therapies that impair fork remodeling or HR, offering potential strategies to selectively eliminate tumor cells with high levels of replication stress. Going forward, it will be important to assess the generality of PPM1D’s role in fork restart across different cell types and biological contexts. Of note, the cell lines used in this study, such as U2OS and HCT116, carry hyper-stabilized forms of PPM1D. It is possible that the dependency on PPM1D varies with the level of ATM signaling, with cells exhibiting higher ATM activity relying more heavily on PPM1D for efficient fork restart, thus exerting selective pressure for mutations that stabilize or increase PPM1D expression.

In conclusion, this study defines a function for the ATM–PPM1D signaling axis in regulating replication fork restart. By limiting ATM activity at stalled forks, PPM1D enables RAD51 recruitment and HR-mediated fork restart and recovery from replication stress. These findings underscore the importance of dynamic phosphorylation control at replication forks and establish PPM1D as a central player in replication stress response pathways with broad relevance to cancer biology and therapeutic intervention.

## METHODS

### Cell lines and cell culture

All cells were maintained at 37 °C with 5% CO_2_ in a humidified incubator and cultured in Dulbecco’s Modified Eagle’s Medium (DMEM) supplemented with 10% bovine calf serum (BCS), 1% non-essential amino acids, 100 U/mL penicillin, and 100 μg/mL streptomycin. U2OS and HCT116 cells were obtained from the American Type Culture Collection (ATCC). Parental U2OS and U2OS-PPM1D-KO cells were kindly provided by Macurek lab. Parental U2OS and U2OS-SMARCAL1-KO cells were kindly provided by Vindigni lab. Parental U2OS and U2OS-53BP1-KO cells were kindly provided by Durocher lab. All cell lines were regularly tested for mycoplasma contamination using the Universal Mycoplasma Detection Kit (ATCC).

### Drugs and reagents

Hydroxyurea (HU, Sigma-Aldrich, #H8627), PPM1D inhibitor (GSK2830371, Sigma-Aldrich, #SML1048), CDC7 inhibitor (XL413, Selleck Chemicals, #S7547), DNA2 inhibitor (C5, MCE, #HY-128729), MRE11 inhibitor (Mirin, Selleck Chemicals, #S8096), ATM inhibitor (AZD0156, MedChemExpress, #HY-100016), ATR inhibitor (AZD6738, MedChemExpress, #HY-19323), and RAD51 inhibitor (B02, Selleck Chemicals, #S8434) were used as indicated in the experiments. Sodium L-ascorbate (Sigma-Aldrich, #A7631) and azide-fluor 488 (Sigma-Aldrich, #760765) were used for the EdU incorporation assay. Dharmacon siGENOME non-targeting control siRNA (siCTL, Horizon Discovery, #D-001210-02-05) and siGENOME human PPM1D (siPPM1D, Horizon Discovery, #D-004554-01-0002) were used for siRNA transfection. 5-Bromo-2’-deoxyuridine (BrdU, Sigma-Aldrich, #10280879001), 5*-*Iodo-2’-deoxyuridine (IdU, Sigma-Aldrich, #I7125), and 5-Chloro-2’-deoxyuridine (CldU, Sigma-Aldrich, #C6891) were used for DNA fiber and combing assays. 2-morpholinoethanesulfonic acid (MES, GoldBio, #M095250), N-lauroylsarcosine sodium salt (Sarkoysl, Sigma-Aldrich, #L5125), and ß-agarase (NEB #M0392L) were used for DNA combing assay. Propidium Iodide (PI, Thermo Fisher Scientific, #P3566) and antifade mounting media with DAPI (Vectashield, #H-1200-10) were used for flow cytometry and imaging experiments as specified.

### Antibodies

Primary antibodies include: β-actin (Proteintech, #66009-1-lg), β-tubulin (Proteintech, #66240-1-lg), Histone H4 (Bethyl Laboratories, #A300-647A), KAP1-pS824 (Bethyl Laboratories, #304-146A), Gamma-H2AX (Bethyl Laboratories, #A300-081A), RPA32 (Bethyl Laboratories, #A300-244A), PCNA (Santa Cruz Biotechnology, #sc-56), CHK1-pS345 (133D3, Cell Signaling Technology, #2348), PPM1D/WIP1 (Santa Cruz Biotechnology, #sc-376257), ATM-pS1981 (Cell Signaling Technology, #4526), 53BP1 (Cell Signaling Technology #4937), RAD51 (Millipore Sigma, #PC130), Rat anti-BrdU (BU1/75 (ICR1) (Abcam, #6326), Mouse anti-BrdU (B44) (BD Biosciences, #347580). Secondary antibodies include: HRP-conjugated goat anti-mouse IgG (Invitrogen #31430), HRP-conjugated donkey anti-rabbit IgG (Cytiva, #NA934V), Alex Fluor 594 goat anti-mouse IgG (H+L) (Invitrogen, #A11005), Alexa Fluor 555 F(ab’)2-Goat anti-rabbit IgG (H+L) (Invitrogen, #A21430), Alexa Fluor 488 goat anti-rabbit IgG (H+L) (Invitrogen, #A11008), Alexa Fluor 488 goat anti-mouse IgG (H+L) (Invitrogen, #A11001).

### Immunofluorescence

For immunofluorescence experiments, cells (25,000 per well) were seeded onto coverslips in 12-well plates. The following day, cells were treated as indicated. Cells were then washed twice with 1 mL ice-cold 1x PBS. Cells were pre-extracted with ice-cold buffer (25 mM HEPES, pH 7.5, 50 mM NaCl, 1 mM EDTA, 3 mM MgCl_2_, 300 mM sucrose, 0.5% Triton X-100) for 5 minutes, followed by a PBS wash. Fixation was performed with 4% paraformaldehyde in PBS for 10 minutes at room temperature (RT), followed by three 5-minute washes in PBS. Cells were permeabilized with 0.25% Triton X-100 in PBS for 5 minutes and washed three times in PBS. Cells were blocked with 5% BSA in 1x PBS for 1 hour at RT. Coverslips were then incubated with primary antibodies (1:300 dilution in 5% BSA) for 1 hour at RT or overnight at 4°C. After three washes with 1x PBS, cells were incubated with fluorophore-conjugated secondary antibodies (1:500 dilution in 5% BSA) for 1 hour at RT in the dark. Following three final PBS washes, coverslips were mounted with DAPI-containing antifade mounting medium, sealed with nail polish, and stored at 4°C. Images were acquired using a Leica DMi8 inverted fluorescence microscope and analyzed using Fiji/ImageJ. Statistical analyses were performed using GraphPad Prism v9.

### EdU incorporation assay

For the EdU incorporation experiment, U2OS cells (25,000 cells per well) were seeded onto coverslips in 12-well plates. The following day, cells were treated as indicated and then processed until permeabilization as immunofluorescence assay. Then, cells were washed by 3% BSA in 1x PBS twice at room temperature (RT). Click reactions were performed with pre-mixed reagents (10 mM Na-L-Ascorbate; 0.1 mM Alexa Fluor Azide; 5 mM CuSO_4_) for 30 minutes at RT. Cells were washed by 3% BSA in 1x PBS twice at RT. Cells were blocked and incubated with primary and secondary antibodies as indicated in the immunofluorescence assay. Following three final 1x PBS washes, coverslips were mounted with a DAPI-containing antifade mounting medium, sealed with nail polish, and stored at 4°C. Images were acquired using a Leica DMi8 inverted fluorescence microscope and analyzed using Fiji/ImageJ. Statistical analyses were performed using GraphPad Prism v9.

### DNA fiber assay

For DNA fiber experiments to monitor replication fork restart, cells were first labeled with 25 μM CldU for the indicated time, then treated with 4 mM HU for 30 minutes to induce fork stalling. After two washes with prewarmed media, cells were incubated in fresh warm media containing 250 μM IdU and 10 µM PPM1D inhibitor, with or without additional inhibitors, for the indicated time. For the fork protection assay, cells were sequentially labeled with 25 μM CldU for 30 minutes and 250 μM IdU for 30 minutes, followed by treatment with 2 mM HU and/or 10 μM PPM1Di or 100 nM ATMi for 90 minutes. Following treatment, cells were harvested by trypsinization, washed, and resuspended in 1x PBS. Approximately 1,000 cells were pipetted onto clean glass microscope slides and lysed with DNA fiber lysis buffer (200 mM Tris-HCl at pH 7.4, 50 mM EDTA, 0.5% SDS) for 2 minutes. Slides were immediately tilted at a 15° angle to stretch DNA fibers, air-dried for 5 minutes, and fixed in a 3:1 methanol/acetic acid solution for 10 minutes at RT. DNA fibers were denatured in 2.5 M HCl for 1 hour, neutralized with 0.4 M Tris-HCl (pH 7.4) for 5 minutes, and washed three times in 1xPBS for 5 minutes each. Slides were blocked for 1 hour in PBS containing 10% goat serum and 10% BSA. Slides were then incubated with mouse anti-BrdU (BD Biosciences, #347580, 1:50) and rat anti-BrdU (Abcam, #ab6326, 1:200) antibodies in PBS with 3% BSA for 1 hour at 37 °C in a humidified chamber. After three 5-min PBS washes, slides were incubated with Alexa Fluor 488-conjugated anti-mouse and Alexa Fluor 594-conjugated anti-rat secondary antibodies (1:500) for 1 hour at RT. Slides were further washed three times in PBS, mounted with Fluoromount-G (Thermo Fisher Scientific), and sealed with nail polish. Images were acquired using a Leica DMi8 inverted fluorescence microscope and analyzed with Fiji/ImageJ software. For fork restart analysis, ongoing forks, bi-directional forks, and terminating forks are classified as restarted forks. The percentages of stalled forks, restarted forks, and newly fired forks were calculated accordingly. Representative images of the different fork types are shown in Fig. S1e. Statistical analyses were performed using GraphPad Prism v9.

### Western blotting

Cells were lysed in ice-cold modified RIPA buffer (50 mM Tris-HCl, pH 7.5; 150 mM NaCl; 1% Tergitol; 0.25% sodium deoxycholate; 5 mM EDTA) supplemented with complete EDTA-free protease inhibitor cocktail (Roche), 1 mM PMSF, and 5 mM NaF. Lysis was performed for 30 minutes at 4 °C and sonicated. Protein concentration was determined using the Bradford assay (Bio-Rad), and 10-15 μg of total protein was resolved by SDS-PAGE. Proteins were transferred to polyvinylidene difluoride (PVDF) membranes and blocked with 5% non-fat milk in TBST (Tris-buffered saline with 0.05% Tween-20) for 1 hour at RT. Membranes were incubated with primary antibodies overnight at 4 °C. The following day, membranes were washed three times with TBST and incubated with HRP-conjugated secondary antibodies for 1 hour at RT. After three additional washes with TBST, membranes were developed using Clarity Western ECL substrate (Bio-Rad, #1705060). Chemiluminescent signals were acquired using a ChemiDoc Imaging System (Bio-Rad) and analyzed with ImageLab software (Bio-Rad).

### Chromatin fractionation

Cells were harvested by trypsinization, washed with PBS, and resuspended in Solution A (10 mM HEPES, pH 7.9; 10 mM KCl; 1.5 mM MgCl_2_; 0.34 M sucrose; 10% glycerol; 0.1% Triton X-100; 1 mM DTT; 10 mM NaF; 1x protease inhibitor cocktail (ThermoFisher, #A32965); 1x PMSF). Cells were incubated on ice for 8 min, then centrifuged at 1,300 x g for 5 minutes at 4 °C. The supernatant was harvested as the cytosolic fraction. The pellet was washed once with solution A, then resuspended in solution B (3 mM EDTA; 0.2 mM EGTA; 1 mM DTT; 1x protease inhibitor cocktail) and incubated on ice for 30 minutes. After centrifugation at 1,700 x g for 5 minutes at 4 °C, the supernatant was collected as the nucleoplasmic fraction. The remaining pellet was washed once with Solution B and resuspended in the 2x sample buffer for analysis as chromatin fraction. Next, chromatin-associated proteins were analyzed by Western blotting.

### Clonogenic assay

U2OS cells (800 cells per well) were seeded in 6-well plates and treated with the indicated drugs for 10 days. At the end of the experiment period, cells were washed once with 1x PBS, fixed in 3.7% paraformaldehyde for 10 minutes at RT, and stained with 0.01% crystal violet in 1x PBS solution. Plates were air-dried and scanned, and colony areas were quantified using ImageJ software.

### Neutral comet assay

Cells were treated as indicated and harvested by trypsinization, followed by a wash with ice-cold 1x PBS. Cells were resuspended at a concentration of 1×10^5^/mL and mixed with molten CometAssay^®^ LMAgarose (R&D Systems, #4250-050-02) at a 1:10 (v/v) ratio. A 50 μL aliquot of the mixture was immediately applied onto CometSlide^®^ (R&D Systems, #4250-050-03) and allowed to solidify at 4 °C for 10 min. Slides were then immersed in lysis solution (R&D Systems, #4250-050-01) for 1 hour at 4 °C. Following lysis, electrophoresis was carried out under neutral conditions at 1V/cm for 30 minutes at 4 °C in the dark. Slides were then incubated in DNA precipitation solution (1M ammonium acetate in 82% ethanol) for 30 minutes at RT, followed by a 30-minute incubation in 70% ethanol. Slides were air-dried at 37 °C and stained with SYBR^TM^ Gold (Thermo Fisher Scientific, #S11494). Images were captured at 20x magnification using a Leica DMi8 inverted fluorescence microscope. DNA damage was assessed by measuring the comet tail moment using OpenComet software (v1.3.1).

### Sample preparation for phosphoproteomic analysis

HCT116 cells were cultured for at least five passages in either “light” or “heavy” SILAC DMEM medium (Thermo Fisher Scientific, #88364) supplemented with 10% dialyzed FBS (Sigma-Aldrich, #F0392), 100 U/mL penicillin, and 100 μg/mL streptomycin (Gibco, #15140-122). Labeling conditions were inverted in biological replicates to filter out phosphopeptides influenced by isotopic bias. After experimental treatments, cells were washed with ice-cold 1xPBS and harvested using a cell scraper. Cell pellets were collected by centrifugation at 1,000 x g for 5 minutes at 4 °C, washed once in PBS, and stored at -80 °C overnight. Frozen pellets were thawed and resuspended in a hypotonic buffer (20 mM Tris-HCl, pH 7.4; 10 mM NaCl; 3 mM MgCl_2_; 0.5 mM DTT) and incubated on ice for 20 min. Cells were centrifuged at 4,500 x g for 4 minutes at 4 °C and the supernatant was discarded. Pellets were resuspended in lysis buffer (50 mM Tris-HCl, pH 8.0; 0.2% Tergitol; 150 mM NaCl; 5 mM EDTA) supplemented with protease inhibitors (ThermoFisher, #A32965), 1 mM PMSF, and 1x PhosSTOP phosphatase inhibitor cocktail (Roche). Lysis proceeded on ice for 30 min, followed by sonication using a Branson 450 digital sonifier with a 102-C probe (15% amplitude, 3s on/ 2s off pulses) for 50 seconds. Lysates were clarified by centrifugation at 45,000 x g for 30 minutes at 4°C. Protein concentrations were determined using the Bradford assay (BioRad). Equal amounts (2 mg each) of light- and heavy-labeled lysates were combined. Samples were denatured and reduced with 1% SDS and 5 mM DTT at 65 °C for 10 minutes. Cysteines were alkylated with 25 mM iodoacetamide for 15 min in the dark. Proteins were precipitated by adding 3 volumes of cold precipitation solution (50% acetone, 49.9% ethanol, 0.1% acetic acid) and incubated on ice for at least 20 minutes. Pellets were recovered by centrifugation at 4,500 x g for 5 minutes at 4°C, washed once with 8 M urea solution and deionized water, and centrifuged at 13,000 x g for 1 minute. Residual water was carefully removed without disturbing the pellet. Pellets were resuspended in 400 µL of 8 M urea in 100 mM Tris-HCl (pH 8.0), then diluted to 2 M urea by adding 1,200 µL of Tris/NaCl solution (50 mM Tris-HCl, pH 8.0; 150 mM NaCl). Proteins were digested overnight at 37 °C with 4 µg of TPCK-treated trypsin. Peptides were acidified to a final concentration 0.2% trifluoroacetic acid (TFA) and formic acid (FA), and desalted using a 200 mg Sep-Pak C18 cartridge (Waters). Briefly, the C18 column was pre-conditioned with buffer B (80% acetonitrile, 0.1% TFA) followed by buffer A (0.1% TFA). Samples were loaded, washed with buffer C (0.1% acetic acid), and eluted with buffer D (80% acetonitrile, 0.1% acetic acid), then dried in a SpeedVac.

Phosphopeptides were enriched using the High-Select Fe-NTA phosphopeptide enrichment kit (Thermo Fisher Scientific, #A32992) following the manufacturer’s protocol. Eluted phosphopeptides were immediately dried and resuspended in 0.1% TFA, followed by a clean-up using a small-scale C18 spin column (Thermo Fisher Scientific, #89879) packed with C18 slurry. Columns were conditioned with buffer B and A, samples loaded, washed with 3% acetonitrile in buffer A, and eluted with buffer D. Eluted peptides were dried in a SpeedVac.

Dried phosphopeptides were resuspended in 15 µL LC-grade H_2_O, 10 µL 10% FA, and 60 µL HPLC-grade acetonitrile. A total of 80 µL was injected into an UltiMate 3000 system and fractionated by HILIC (hydrophilic interaction liquid chromatography) using a TSKgel Amide-80 column (2 mm × 150 mm, 5 μm; Tosoh Bioscience). The following gradient buffers were used: buffer A (90% acetonitrile), buffer B (75%acetonitrile and 0.005% TFA), and buffer C (0.025% TFA). The gradient program was as follows: 100% buffer A (0 min); 94% buffer B/6% buffer C (3 min); 65.6% buffer B/34.4% buffer C (30 minutes); and 5% buffer B/95% buffer C (32 - 40 minutes). Flow rate: 150 µL/min. Fractions were collected every 30 seconds from minute 5 to minute 24. All fractions were dried, resuspended in 5 μL of 0.1P (1 pmole/µL angiotensin in 0.1% TFA), and subjected to LC-MS/MS analysis.

### ProKAS biosensor sample preparation

Stable U2OS cell lines expressing the ProKAS construct were generated using lentiviral transduction, followed by selection with 1 µg/mL puromycin. Following treatment, cells cultured in 6-well tissue culture plates were lysed using a modified radioimmunoprecipitation assay buffer (mRIPA) buffer supplemented with 0.2% SDS (SDS lysis buffer). After aspirating the culture media, 175 µL of SDS lysis buffer was added to each well. Plates were placed on a hot plate set to 200°C for 10 seconds and immediately removed and agitated to promote the lysis. This heating and agitation step was repeated 2-3 times to ensure complete lysis. Following lysis, 525 µL of mRIPA buffer was added to each well to dilute the final SDS concentration to below 0.1%. Lysates were collected into pre-labeled tubes and sonicated using a Branson 1/8” probe tip sonifier (5 seconds at 12% amplitude) to shear genomic DNA and reduce viscosity. Samples were then centrifuged at 16,000 x g for 3 minutes at 4°C to pellet debris. For affinity purification, 4 µL of ALFA magnetic resin (NanoTag, #N1516) was used per lysate. Resin was equilibrated by resuspending in 1 mL mRIPA buffer, and then distributed evenly to LoBind Eppendorf tubes placed on magnetic racks. Afterwards, the buffer was aspirated, and the lysates were transferred to the tubes containing the immobilized resin. The samples were incubated for 30 minutes at 4°C on a nutating mixer. After incubation, beads were immobilized on magnetic racks and supernatants were aspirated. Beads were washed twice with 500 µL mRIPA buffer and once with 1 mL sterile deionized water, with each wash involving nutation for 1 minute at RT. The beads were then immobilized and 80 µL of acidic elution buffer (0.1 M Glycine-HCl, 150 mM NaCl, pH 2.2) was added to each tube. The tubes were agitated for 2 minutes at RT to aid in elution of the bait protein. Eluted material was collected into new LoBind Eppendorf tubes. For digestion, each elution was adjusted by adding 20 µL of 1M Tris (pH 8.0), 50 µL of 8 M Urea, and 50 µL of 150 mM NaCl-50 mM Tris (pH 8.0) containing 100 ng Trypsin Gold (Promega, #V5280). Samples were incubated at 37 °C on a nutating mixer for 6-18 hours to allow tryptic digestion of immunoprecipitated proteins. Peptide samples were acidified by adding 10 µL of 10% Trifluoroacetic acid (TFA) and 10 µL of 10% Formic Acid. Acidified digests were desalted using 20 mg of C18 resin (Sep-Pak cartridge; Waters, WAT054945) packed into Pierce Micro Spin Columns (ThermoFisher Scientific, #89879). The resin was conditioned with 200 µL of 80% acetonitrile (ACN) containing 0.1% TFA, followed by 200 µL 0.1% TFA. Acidified samples were then loaded onto the resin, washed with 350 µL of 0.1% acetic acid, and peptides were eluted into LoBind Eppendorf tubes using 100 µL of 80% ACN containing 0.1% acetic acid. Eluted peptides were then dried in a vacuum concentrator and immediately resuspended in 10 µL of 250 mM HEPES (pH 8.0) for tandem mass tag (TMT) labeling. A 50 µg aliquot of TMTsixplex reagent (ThermoFisher Scientific, #90061) was used to label up to 6 different sample conditions. For each TMT channel, 2 µL of ACN was added to the reagent, followed by the immediate addition of 8 µL of resuspended peptide. Labeling reactions were incubated for 60 minutes at RT and quenched by adding 2 µL of 5% hydroxylamine. Subsequently, 25 µL of 0.1% TFA was added to each channel. Samples from all channels were then pooled, and an additional 30 µL of 10% TFA was added to the combined sample. The pooled TMT-labeled peptides were desalted again using C18 cleanup as described above. The dried samples were resuspended in LC-grade water prior to MS injection.

### DNA combing assay

For the DNA combing assay, 200,000 cells were seeded and cultured for 24 hours before labeling. Cells were first labeled with 25 µM CldU for 20 minutes, followed by treatment with 4 mM HU for 30 minutes. Cells were washed twice with pre-warmed 1x PBS and incubated in pre-warmed complete medium containing 250 µM IdU, with or without 10 µM PPM1Di for 40 minutes. Cells were harvested and resuspended in 1.2 % low-melting-temperature (LMT) agarose (Sigma-Aldrich, #A4018) pre-warmed to 55 °C and allowed to solidify at 4 °C for 1 hour. Cell-containing agarose was incubated in ESP solution (200 µL 0.5 M EDTA, pH 8.0; 25 µL 10% Sarkosyl; 50 µL Proteinase K) at 50 °C for 16 hours. The ESP solution was then discarded, and cells were washed six times with a modified TE buffer (10 mM Tris-HCl, 1 mM EDTA, pH 8.0). Agarose plugs were melted in 1 mL MES buffer (1.22 g MES dissolved in 12.5 mL ddH_2_O, pH 5.5) at 68 °C for 20 minutes and further incubated at 42 °C for 20 minutes. Subsequently, 1.0 µL ß-agarase was added to each sample and digestion was performed at 42 °C for 12 hours. DNA solutions were transferred into disposable reservoirs (Genomic Vision) and loaded onto the molecular combing system (Genomic Vision). Functionalized microarray coverslips (PolyAn, #10490804) were used for DNA stretching. After combing, coverslips were baked at 65°C for 2 hours and fixed in methanol for 10 minutes at RT. DNA fibers were denatured, blocked, incubated with primary and secondary antibodies as described for the DNA fiber assay. Images were acquired and analyzed as DNA fiber assay. Representative images of different combing conditions are shown in Fig. 1d.

### siRNA transfection

For siRNA transfection performed in EdU incorporation assay, 10,000 cells per condition were seeded on 6cm plates and incubated at 37 °C until 60%-70% confluent. Then, solution A (3 µL of siRNA (20 pmols) in 250 µL Opti-MEM) and solution B (9 µL of Lipofectamine in 250 µL Opti-MEM) were incubated for 5 minutes at room temperature before mixing. After incubating for 20 minutes, the transfection mixture was added drop-wise on cells. Transfection reagents were removed after 6 hours and cells were cultured with complete medium for at least 48 hours, before being treated as indicated in each experiment.

### MS data acquisition

#### Phosphoproteomics

Phosphopeptide fractions were resuspended in 0.1% trifluoroacetic acid and analyzed by liquid chromatography-tandem mass spectrometry (LC-MS/MS) using an UltiMate 3000 HPLC system coupled to a Q-Exactive HF mass spectrometer (Thermo Fisher Scientific). Peptides were separated on a 35 cm in-house packed C18 column (inner diameter of 100 µM) using reversed-phase chromatography. The Q-Exactive HF was operated in data-dependent acquisition (DDA) mode. Full MS scans were acquired in the Orbitrap over a mass-to-charge range of 380-1800 *m/z* at a resolution of 60,000. The 20 most abundant precursor ions with charge states of +2, +3, and +4 were selected for MS/MS fragmentation using higher-energy collisional dissociation (HCD) with a normalized collision energy (NCE) of 28. Fragment ion spectra were acquired in the Orbitrap at a resolution of 15,000 with an AGC target of 1×10^5^ and a maximum injection time of 100 ms. Dynamic exclusion was enabled with a 30-second exclusion window to prevent repeated sequencing of previously fragmented ions.

#### ProKAS Sensor

To check the TMT labeling efficiency and retention time of labeled peptides, data-dependent analyses were performed on the Q Exactive HF mass spectrometer using a 70-minute method containing a 60,000 scanning resolution, AGC target of 3×10^6^, maximum inject time of 30 ms, and scan range between 380 and 1800 m/z. The MS2 parameters included a 15,000 resolution, AGC target of 1×10^5^, maximum inject time of 100 ms, isolation window of 2.0 m/z and normalized collision energy (NCE) of 36. Once the mass of target peptides and their phosphorylated forms were identified, parallel reaction monitoring was performed utilizing a 70-minute method with 15,000 resolution, AGC of 2×10^5^, an inject time of 120 ms, and an isolation window of 0.4 m/z. Labeling efficiency was determined to be between 85-90%, and parallel reaction monitoring libraries were determined based on Comet searches to design an inclusion list with peptide masses.

### Phosphoproteomic data analysis

Raw LC-MS/MS data were analyzed using the Comet search engine. The following search parameters were applied: precursor mass tolerance of ±10 ppm; semi-tryptic digestion; variable modifications of +8.0142 Da (Lysine, heavy SILAC), +10.00827 Da (Arginine, heavy SILAC), +79.966331 Da (phosphorylation of Ser/Thr/Tyr), and +57.021465 (carbamidomethylation of cysteine). Searches were performed against the Uniprot human proteome database (74,304 entries). Peptide identification was refined using PeptideProphet, with the following parameters: minimum probability of 0.05, minimum peptide length of 7 residues, accurate mass binning, charge states restriction to +2, +3, and +4, and phospho-site scoring enabled (“use Phospho-information”). Peptide quantification was performed using Xpress, with a mass tolerance of 0.005 Da, requiring a minimum of one point for quantitation and one isotopic peak per peptide. Phosphosite localization confidence was determined using PTMProphet. In-house R-scripts were used to further process the data, including: grouping adjacent phosphorylatable residues into phosphosite clusters when exact localization could not be determined; calculating the average and median Xpress ratio for each peptide; computing statistical significance using the Mann-Whitney U test.

### ProKAS Data Analysis with Skyline

To construct the spectral libraries utilized for ProKAS parallel reaction monitoring, raw MS/MS data generated from the DDA runs were searched on the Comet search engine to obtain peptide masses. The search was conducted using the following parameters: semi-tryptic peptide identification, mass tolerance within 15 ppm, variable modifications for STY phosphorylation (79.966331 Da). The search was performed against the Uniprot human proteome database in addition to ProKAS peptide sequences, as well as the TMT condition. PTMprophet was utilized to determine phosphorylation site localization, search results were imported into Skyline, containing masses of the unphosphorylated and phosphorylated forms of the target ProKAS peptides. Once a library was generated, PRM data was imported into Skyline for analysis by conversion to mzXML format using msConvert. Transitions were verified manually before generating a transitions report from Skyline containing the Protein, Peptide Modified Sequence, Replicate, Precursor Mz, Precursor Charge, Product Mz, Fragment Ion, Retention Time, and Area. Data in these reports were processed for each TMT fragment ion in each replicate by normalization of phosphorylated peptide ratios to their unphosphorylated counterparts. Then, the normalized ratios in each condition were compared against the normalized ratios in the untreated condition to calculate changes in phosphorylation levels for each peptide. Each condition was run in three replicates and error bars in associated figures reflect the standard deviation of the replicates in each condition.

### Statistical analysis

Statistical analyses were performed using GraphPad Prism. When comparing three or more groups, statistical significance was assessed using one-way ANOVA. For comparisons between two groups, a two-tailed unpaired Student’s *t*-test was used. Exact *p*-values, statistical tests, and replicate numbers are detailed in the corresponding figure legends.

## Data Availability

Mass spectrometry data generated for this study are available through PRIDE, wherein phosphoproteomics data can be found under PXD identifier PXD045462 (https://www.ebi.ac.uk/pride/)(69).

## ACKNOWLEDGEMENTS

We thank Libor Macurek for providing U2OS WT and PPM1D-KO cells, Daniel Durocher for providing U2OS WT and 53BP1-KO cells, and Alessandro Vindigni for providing U2OS WT and SMARCAL1-KO cells. We acknowledge the Weill Institute for providing access to the ChemiDoc Imaging system used in this study. We thank Catalina Pereira and Munisha Mujingjiang for providing supplies and technical support to perform DNA combing experiments. We thank Beatriz Almeida for technical assistance and all members of the Smolka lab for helpful discussions and technical support. M.B.S. was supported by NIH grant R35-GM563209. Y.W. was supported by a postdoctoral fellowship from the Fleming Research Foundation.

## AUTHOR CONTRIBUTIONS

JB and MBS led the conceptualization of the main study, defining the problem statement and research objectives. YC, YW, and MBS contributed to the conceptualization and refinement of the final study design. JB performed and analyzed immunofluorescence, western blot, chromatin fractionation, comet assays, clonogenic assays, and DNA fiber assays for the initial study. YC performed and analyzed the final experiments for all immunofluorescence, EdU incorporation, western blot, siRNA transfection, and DNA combing assays. YW conducted and analyzed the final experiments for BrdU incorporation, neutral comet assays, and DNA fiber assays. YW and TCH jointly performed and analyzed the clonogenic survival assays. JB and VMF carried out and analyzed the phosphoproteomic experiments. KJ performed and analyzed the ProKAS experiments. EÇ contributed to DNA fiber data analysis for the pilot study. JB, YC, and YW prepared the data visualizations. MBS and JB wrote the initial version of the manuscript, with YC and YW contributing to the final version. YC, YW, KJ, and MBS reviewed and edited the manuscript. MBS secured funding and supervised the project. All authors contributed to and approved the final version of the manuscript.

---

**Figure S1.**
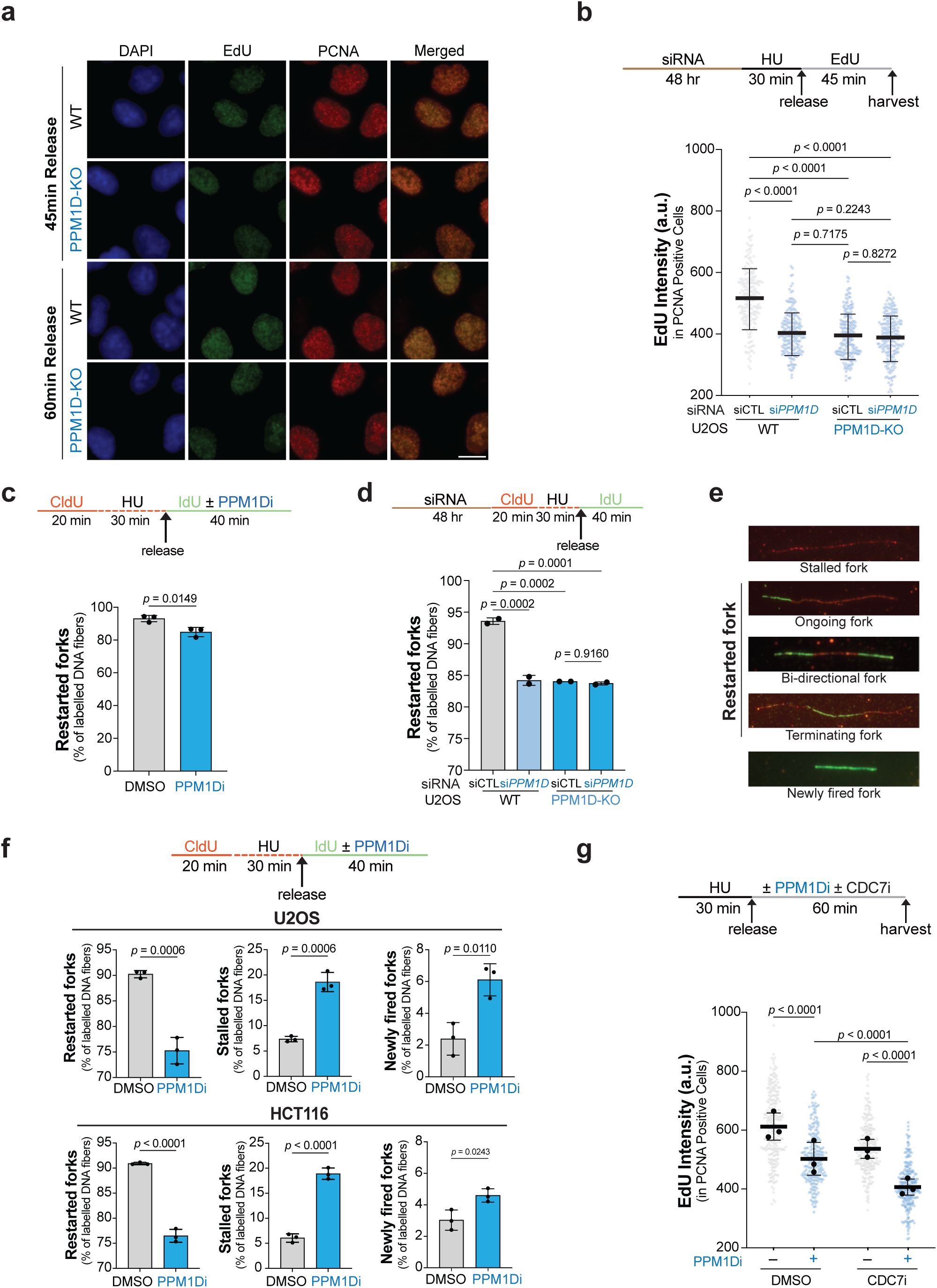
PPM1D modulates DNA replication. (a) Representative immunofluorescence images from the EdU incorporation assay shown in Fig. 1c. Scale bar, 10 μm. (b) EdU incorporation analysis in U2OS wild-type (WT) and PPM1D-knockout (KO) cells transfected with control or PPM1D siRNA (siCTL and si*PPM1D*). Top: experimental scheme. Bottom: dot plot showing EdU intensity in PCNA-positive cells from one representative experiment. A minimum of 400 cells were quantified per condition from two independent biological replicates. *p*-values were determined by one-way ANOVA with Tukey’s multiple comparisons test. Error bars represent standard deviation. (c) DNA combing assay in U2OS cells. Top: experimental scheme. Bottom: bar graph showing the percentages of restarted replication forks in cells treated with or without PPM1Di (10 μM). A minimum of 2,720 fibers were quantified per condition from three independent biological replicates. *p*-values were calculated using Student’s t-test. Error bars represent standard deviation. (d) DNA combing assay in U2OS wild-type (WT) and PPM1D-knockout (KO) cells transfected with control or PPM1D siRNA (siCTL and si*PPM1D*). Top: experimental scheme. Bottom: bar graph showing the percentages of restarted replication forks. A minimum of 1,644 fibers were quantified per condition from two independent biological replicates. *p*-values were calculated using one-way ANOVA with Tukey’s multiple comparisons test. Error bars represent standard deviation. (e) Representative images of different fork types in the DNA fiber assay. (f) DNA fiber assay in U2OS and HCT116 cells. Top: experimental scheme. Bottom: bar graphs show the percentages of restarted, stalled and newly fired replication forks in cells treated with or without PPM1Di (10 μM). A minimum of 471 fibers (U2OS) and 641 fibers (HCT116) were quantified per condition from three independent biological replicates. *p*-values were calculated using Student’s t-test. Error bars represent standard deviation. (g) EdU incorporation analysis in U2OS cells. Top: experimental scheme illustrating treatment with the PPM1D inhibitor (10 µM) and/or CDC7 inhibitor (CDC7i; XL413; 20 µM). Bottom: dot plot showing EdU intensity in PCNA-positive cells pooled from three independent biological replicates (at least 73 cells per condition per replicate). The mean EdU intensity from each replicate is indicated by black dots. *p*-values were determined by one-way ANOVA with Tukey’s multiple comparisons test. Error bars represent standard deviation.

**Figure S2.**
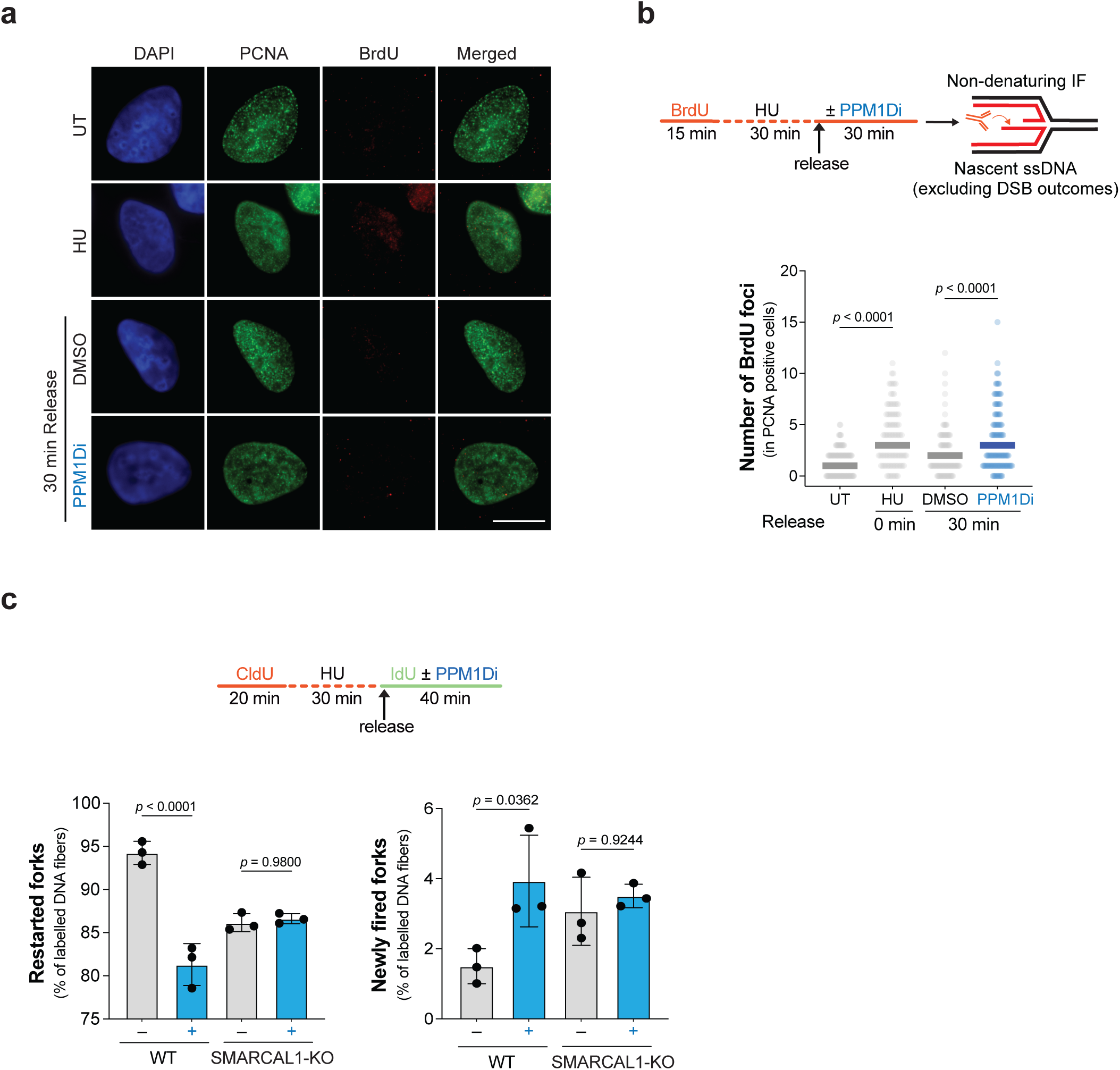
PPM1D regulates RPA32 foci accumulation and replication-associated ssDNA formation. (a) Representative immunofluorescence images corresponding to Fig. 2e. (b) Analysis of nascent single-stranded DNA (ssDNA) in U2OS cells using BrdU immunostaining under non-denaturing conditions, as shown in the treatment scheme. Dot plot shows the number of BrdU foci in PCNA-positive cells from a second biological replicate. At least 113 PCNA-positive cells were quantified per condition. *p*-values were calculated using one-way ANOVA. (c) DNA fiber assay in U2OS WT and SMARCAL1-KO cells. Bar graphs showing the percentage of restarted (left) or newly fired (right) replication forks in the presence or absence of PPM1D inhibitor (PPM1Di). At least 942 fibers per condition were quantified per condition across three independent biological replicates. *p*-values were calculated using one-way ANOVA with Tukey’s multiple comparisons test. Error bars represent standard deviation. A schematic of the fiber assay condition is shown at the top of the panel.

**Figure S3.**
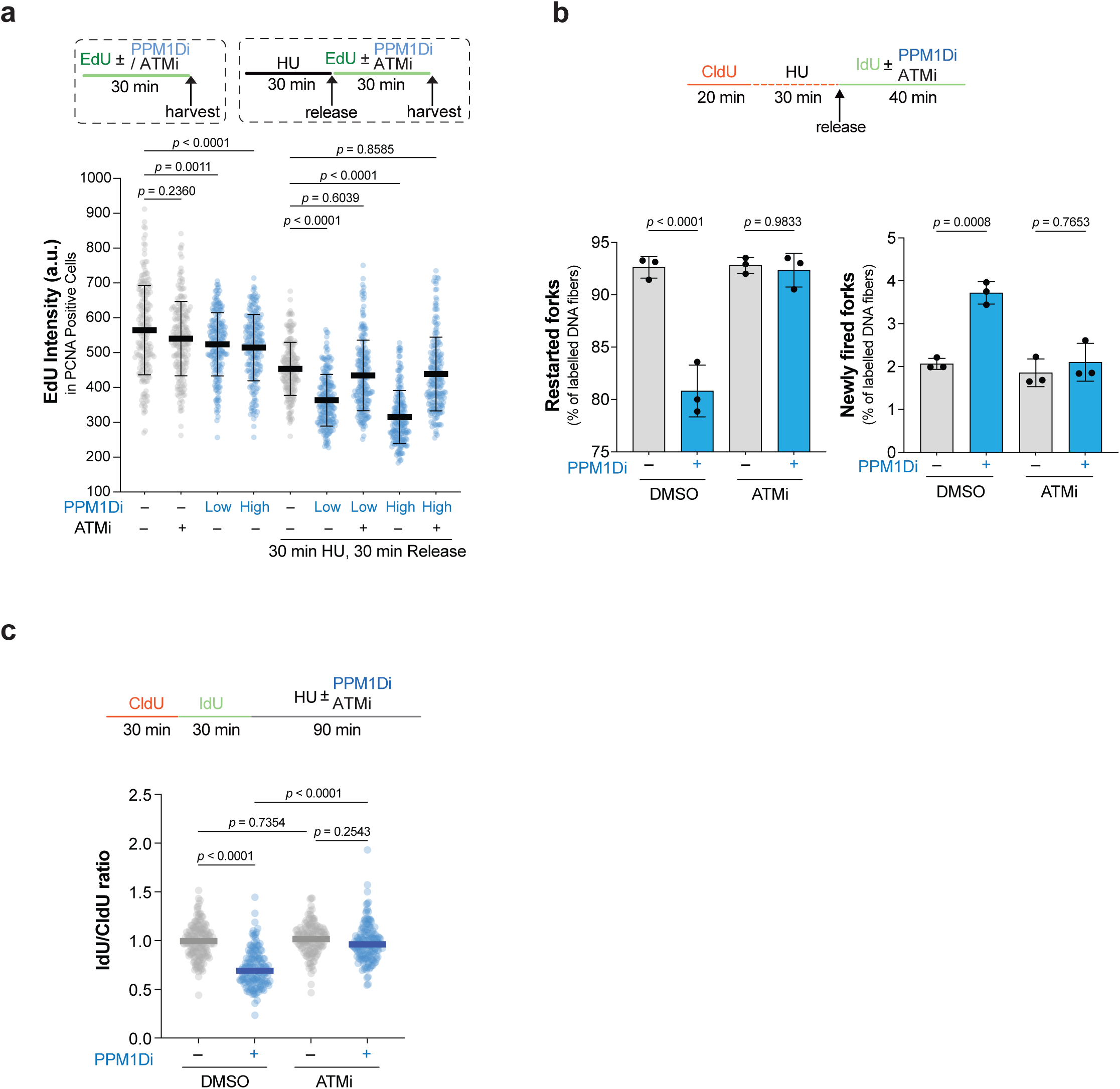
ATM inhibition rescues replication fork restart. (a) EdU incorporation assay in U2OS cells. Top: experimental schemes showing treatment with and without HU (4 mM), PPM1Di (Low dose: 3 µM, High dose: 10 µM) and ATMi (100 nM). Bottom: dot plot shows the EdU intensity in PCNA-positive cells from one representative experiment (at least 196 cells per condition). *p*-values were determined by one-way ANOVA with Tukey’s multiple comparisons test. Error bars represent standard deviation. (b) DNA fiber assay in U2OS cells treated with or without PPM1Di (10 µM) and ATMi (100 nM). Bar graphs showing the percentage of restarted (left) or newly fired (right) replication forks under each treatment condition across three independent biological replicates. *p*-values were calculated using one-way ANOVA with Tukey’s multiple comparisons test. Error bars represent standard deviation. A schematic of the fiber assay condition is shown at the top of the panel. (c) DNA protection assay in U2OS cells treated with or without PPM1Di (10 µM) and ATMi (100 nM), as indicated. Dot plot showing the ratio of IdU/CldU tract lengths as indicator of replication fork degradation from a second biological replicate. At least 144 fibers were analyzed per condition. *p*-values were calculated using one-way ANOVA.

**Figure S4.**
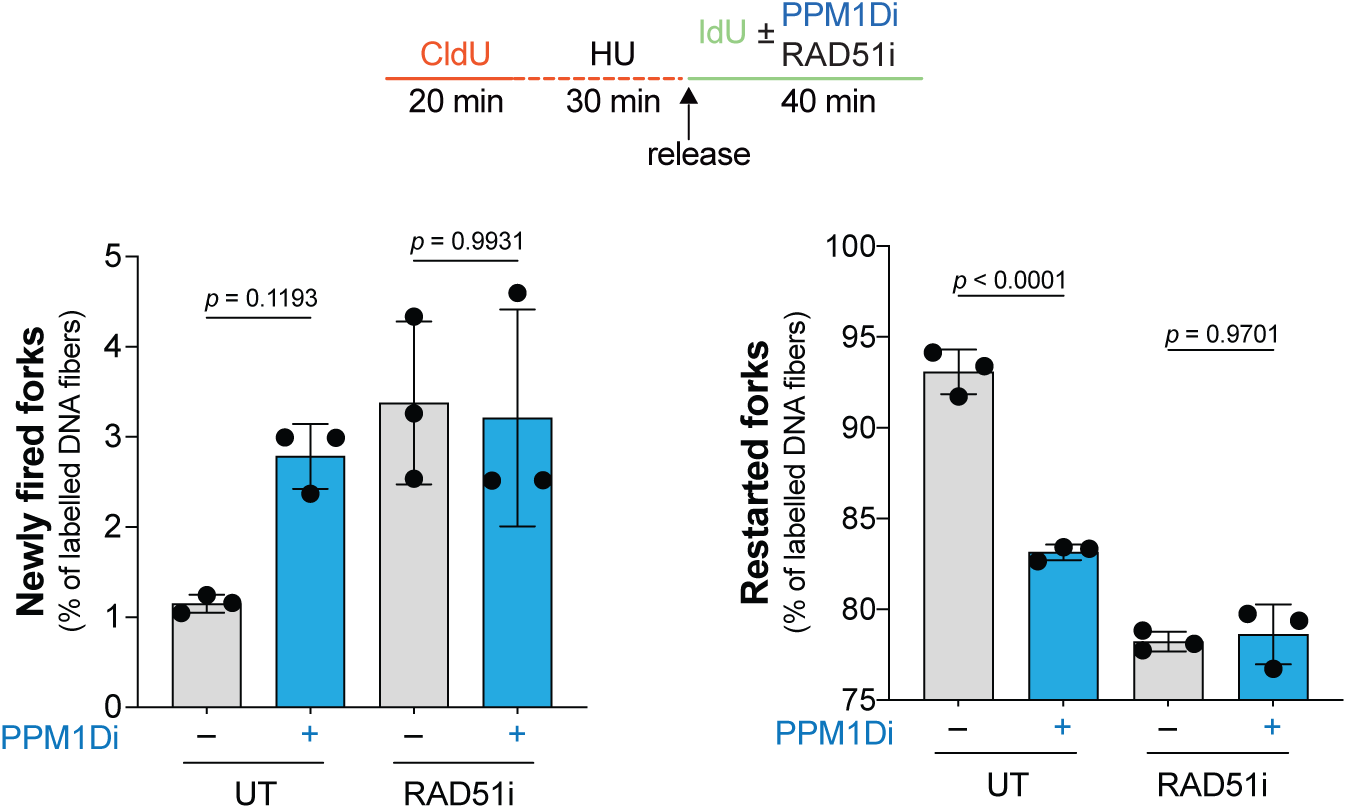
PPM1D regulates RAD51-mediated replication fork restart. DNA fiber assay in U2OS cells treated with or without PPM1Di (10 µM) and RAD51 inhibitor (RAD51i; B02; 20 µM). Bar graphs showing the percentage of newly fired (left) or restarted (right) replication forks under each treatment condition across three independent biological replicates. *p*-values were calculated using one-way ANOVA with Tukey’s multiple comparisons test. Error bars represent standard deviation. A schematic of the fiber assay condition is shown at the top of the panel.

**Figure S5.**
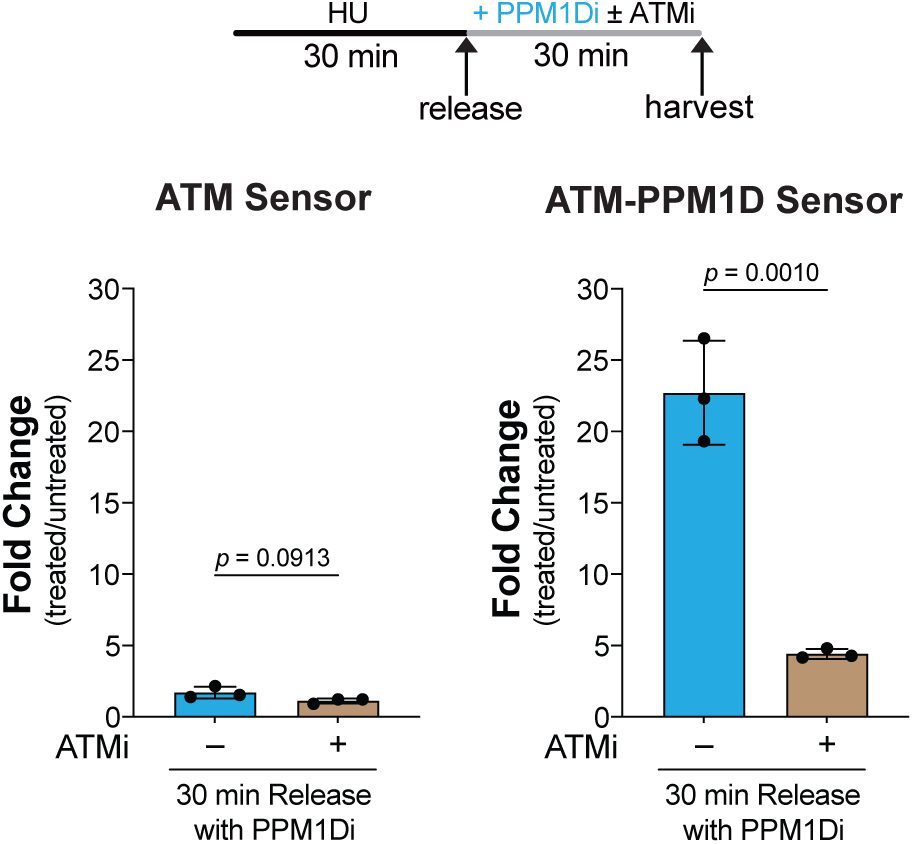
PPM1D inhibition induces phosphorylation of ProKAS sensors in an ATM-dependent manner. Quantification of ATM and PPM1D activity using ProKAS sensors during replication stress recovery in the presence of PPM1Di (10 µM), with or without ATMi (100 nM). Bar graphs represent the normalized ATM activity (left) and PPM1D acitivity (right) relative to untreated condition from three independent biological replicates. *p*-values were calculated using Student’s t-test. Error bars represent standard deviation.

